# Deciphering Oxygen Distribution and Hypoxia Profiles in the Tumor Microenvironment: A Data-Driven Mechanistic Modeling Approach

**DOI:** 10.1101/2024.03.04.583326

**Authors:** P. Kumar, M. Lacroix, P. Dupré, J. Arslan, L. Fenou, B. Orsetti, L. Le Cam, D. Racoceanu, O. Radulescu

## Abstract

The distribution of hypoxia within tissues plays a critical role in tumor diagnosis and prognosis. Recognizing the significance of tumor oxygenation and hypoxia gradients, we introduce mathematical frameworks grounded in mechanistic modeling approaches for their quantitative assessment within a tumor microenvironment. Our approach provides a non-invasive method to measure and predict hypoxia using known blood vasculature. Formulating a reaction-diffusion model for oxygen distribution, we apply it to derive the corresponding hypoxia profile. The modeling and simulations successfully replicate the observed inter- and intra-tumor heterogeneity in experimentally obtained hypoxia profiles across various tumor tissues (breast, ovarian, and pancreatic) in our dataset. Employing a data-driven approach, we propose a method to deduce partial differential equation (PDE) models with spatially dependent parameters, enabling us to comprehend the variability of hypoxia profiles within a tissue. The versatility of our framework lies not only in capturing diverse and dynamic behaviors of tumor oxygenation but also in categorizing states of vascularization. These categories are distinguished based on the dynamics of oxygen molecules, as identified by the model parameters.

## 1 Introduction

Within several solid tumors, increased metabolic demands and abnormal vasculature lead to the development of regions deprived of adequate oxygen availability. This state of tissue microenvironment is termed hypoxia, and consequently, the regions are known as hypoxic [36]. Hypoxia, considered as one of the hallmarks of most solid tumors, compromises the standard functionality of cells and even survival, resulting in the adaptation of (cancer) cells and their dynamics from transcription level to population one. These adaptations affect multiple cellular pathways and contribute to tumor progression, invasiveness, angiogenesis, and vasculogenic mimicry [10]. Furthermore, hypoxia and its mediators, e.g., hypoxia-inducible factor (HIF), influence multiple signaling pathways and gene regulation to promote neovascularization, invasion, migration, adhesion, metastasis, pH regulation, DNA replication, protein synthesis and phenotypic switching (from proliferative to invasive) [55, 1, 18]. Thus, hypoxia-driven changes result in intra-tumor heterogeneity, which hinders accurate diagnosis. Another crucial effect of hypoxia is observed in the reduced treatment efficacy, as the hypoxic regions of the neoplasm do not respond well and show more resistance towards treatment, especially radiation therapy. Such effects are more evident in a few aggressive tumors, such as melanoma, one of the most aggressive, complex, and heterogeneous cancers [4, 17, 10]. The alleviation of hypoxia in a tissue region can be achieved through several strategies; one involves an increment in oxygen delivery, and another focuses on the decrement of oxygen consumption by the cancer cells [13]. The former is a widely accepted procedure [22, 46] and few studies, e.g., [13, 43, 32], suggest that the latter can also be efficient in overcoming hypoxia and its consequences. Hence, accurate assessments of oxygen distribution, hypoxia profile, and *O*_2_ consumption in the tumor microenvironment would be helpful for the prognostic evaluation and therapeutic options.

Mathematical modeling has proven to be beneficial in describing complex dynamics, offering powerful and cost-efficient tools for making predictions in various fields. In the context of tumor dynamics, mathematical models are valuable for reconciling spatiotemporal and phenotypic heterogeneity, as well as designing improved treatment strategies [2, 21]. Numerous computational and mathematical models have been proposed to describe tissue oxygenation, especially focusing on the distribution of oxygen in both cancerous and regular tissues [49, 15, 23, 42, 44, 16].

The cylinder model, initially proposed in [30], laid the foundation for calculating oxygen flow based on an analytical expression dependent on the difference in oxygen tension in steady-state. Subsequently, several partial differential equation (PDE) models of the reaction-diffusion type were developed to represent the oxygen spatial distribution [49, 23, 42, 44, 45, 38, 54, 11]. These models involve different forms of source terms and boundary conditions. The resulting PDEs were numerically solved through discretization methods (finite-difference or finite element) or numerical integration, as seen in the alternative approach based on Green’s function method [23, 16]. This method was later modified in [42, 44] to account for more complex vascular architecture.

Despite these advancements, only a few works explicitly discuss hypoxia in connection with real-world tumor samples or experimental data. For instance, in [38], the predicted hypoxia, observed using the needle electrode technique, is qualitatively compared with CD31^1^ stained images. However, most formulations are limited to synthetic data and qualitative results, lacking direct application and connection with actual tumor tissues and experimental outcomes. Hence, there is still a need for accurate predictive models of hypoxia that take into account tissue heterogeneity.

Here, we propose a physical model that predicts hypoxia in heterogeneous tumor microenvironments from stained tissue images. For this aim, we develop a data-driven mathematical framework to model the distribution of oxygen and hypoxia in the tumor microenvironment, incorporating mechanistic inferences. We produce experimental data in the form of high-pixel images that contain staining of tumor tissues, serving as a proxy for blood vessels, hypoxia, and cell nuclei. The identification of these three elements is detailed as follows:

- Blood Vessels: The CD31 marker is a proxy for blood vessels. CD31, also known as PECAM-1 (platelet endothelial cell adhesion molecule 1), is a 130-kDa transmembrane glycoprotein belonging to the immunoglobulin superfamily. It is present on the surface of platelets, monocytes, macrophages, and neutrophils and is a constituent of the endothelial intercellular junction [39]. Endothelial cells form a continuous lining (endothelium) along the inner surface of blood vessels, providing a barrier between circulating blood and surrounding tissues. By identifying these cells, the desired blood vessels are marked.
- Hypoxia: Intra-tumoral hypoxia is responsible for activating the key transcription factor HIF, which mediates the activation of several defense mechanisms for tumor cells. HIF-1 is a heterodimeric transcription factor comprising an HIF-1*α* subunit and an HIF-1*β* subunit [33]. HIF-1, promoted by intra-tumoral hypoxia, regulates CA9. CA9 is considered one of the best-characterized targets of HIF-1 [5], serving as an intrinsic marker for tumor hypoxia^2^.
- Cell Nuclei: The DAPI marker, which stands for 4’,6-diamidino-2-phenylindole, is used to identify cell nuclei [47, 26]. DAPI is a standard marker for detecting nuclei.

The acquired images exhibit intra- and inter-tumor heterogeneity in oxygen and hypoxia distribution. Leveraging this dataset, we develop a mechanistic framework to model oxygen and hypoxia distribution, predicting the hypoxia profile for a tumor tissue based on known vasculature.

Our study successfully recovers the biologically relevant heterogeneity of oxygenation and identifies crucial parameters involved in reducing hypoxia. As hypoxia promotes radioresistance [53, 32], these results are also valuable for enhancing the effectiveness of radiotherapy. We modeled oxygen distribution using a steady-state solution of a reaction-diffusion PDE. For a known distribution of *O*_2_, we modeled the hypoxia profile based on the dynamics of a proxy for HIF, namely the carbonic anhydrase IX (CA9) biomarker. While similar theoretical and computational models for oxygen distribution exist in the literature [6, 49, 28], the novelty of our work lies in identifying and validating a predictive theoretical model using tumor samples and addressing the challenge of hypoxia heterogeneity. More precisely, we demonstrate the derivation of a space-dependent coefficient partial differential equation (PDE) model that predicts oxygen and hypoxia gradients. This is achieved by starting from a static model with space-invariant coefficients and identifying proxies for space-dependent coefficients directly from images of stained tissues.

The remainder of the article is organized as follows. The methods used in this work are detailed in Section 2, covering information about the experimentally produced data, data processing, quantification of blood vessel diversity, mathematical modeling of oxygen and hypoxia, numerical simulation, parameter estimation, and validation. The obtained results are discussed in Section 3. The study is concluded with a discussion in Section 4 and a summary in Section 5.

## 2 Methods

### 2.1 Experimental Data

#### 2.1.1 Primary derived xenograft (PDX)

A single fragment of frozen tumor, approximately 8 mm^3^, is implanted into the inter-scapular fat pads of 3 to 4-week-old female Swiss-nude mice. Tumor growth is measured weekly using calipers, aiming for a maximum of 1500 mm^3^ [9, 34]. Tissues from three cancer types are engrafted: breast (Triple Negative Breast Cancer #0056, #0520, #1090; Her2+ Breast Cancer #1092), ovarian (High-Grade Serous Ovarian Cancer #0721), and pancreatic (#1087, #1137, #1905) as listed in table 1.

**Table 1:**
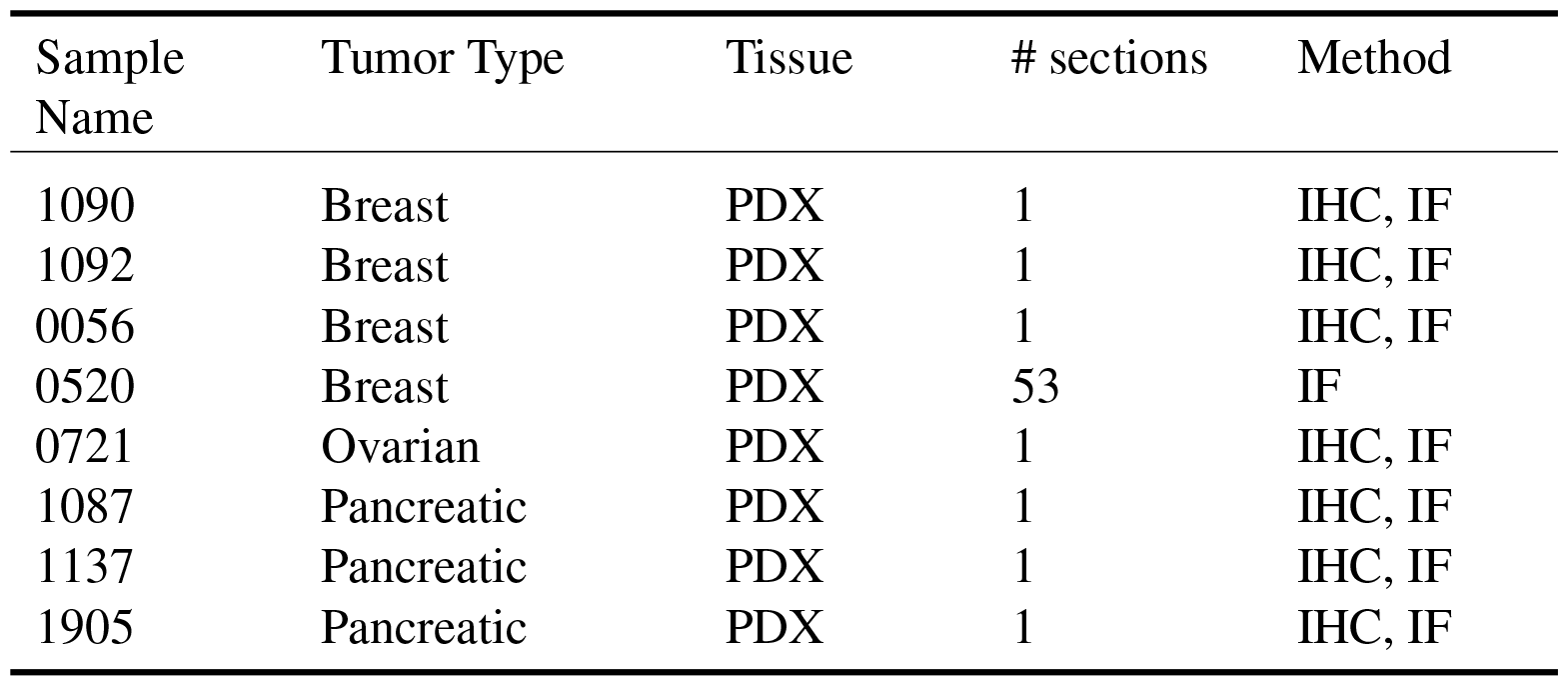
Produced experimental data: In this work, we perform staining on three different PDX tumor types, namely breast, ovarian, and pancreatic. This table lists these tumor samples, with the first column containing the name of the sample used during the experiment. For each PDX, both staining techniques are applied to different tissue samples, generating one stained for IF and one for each CD31 and CA9 with IHC scans. Except for the tumor 0520, where IF is performed on multiple tissues of that tumor sample.

#### 2.1.2 Immunohistochemistry and Immunofluorescence

PDX samples are fixed in 4% neutral-buffered formalin for 24 hours and then paraffin embedded. The FFPE tissues are sectioned into 4*µ*m sections and processed for immunohistochemistry (IHC) or Immunofluorescence (IF). CD31 (Abcam, #ab28364) and CA9 (Roche, #760-6080) stainings are carried out on a VENTANA Discovery Ultra automated staining instrument (Ventana Medical Systems) following the manufacturer’s instructions. For IF, after applying the primary antibodies, sections are incubated for 1 hour at room temperature with secondary antibodies coupled to Alexa-488, Alexa-647 (Life Technologies). Nuclei are counterstained with 4’,6-diamidino-2-phenylindole (DAPI, Sigma), washed in PBST, and coverslipped using Mowiol mounting solution. For image analyses, sections are scanned on a 3DHISTECH scanner.

In IHC, stainings for CA9 and CD31 are conducted on adjacent tissue sections spaced four microns apart. The outcome reveals a brown color, indicating the presence of the targeted proteins, accompanied by cell nuclei depicted in blue, as exemplified in fig. 1 (middle and right columns). Conversely, IF stainings are performed on the same tissue section with different target proteins represented by specific colors. Specifically, CD31, CA9, and DAPI are shown in red, green, and blue colors, respectively. The left column images in fig. 1 visually demonstrate these stains in the mentioned colors.

**Figure 1:**
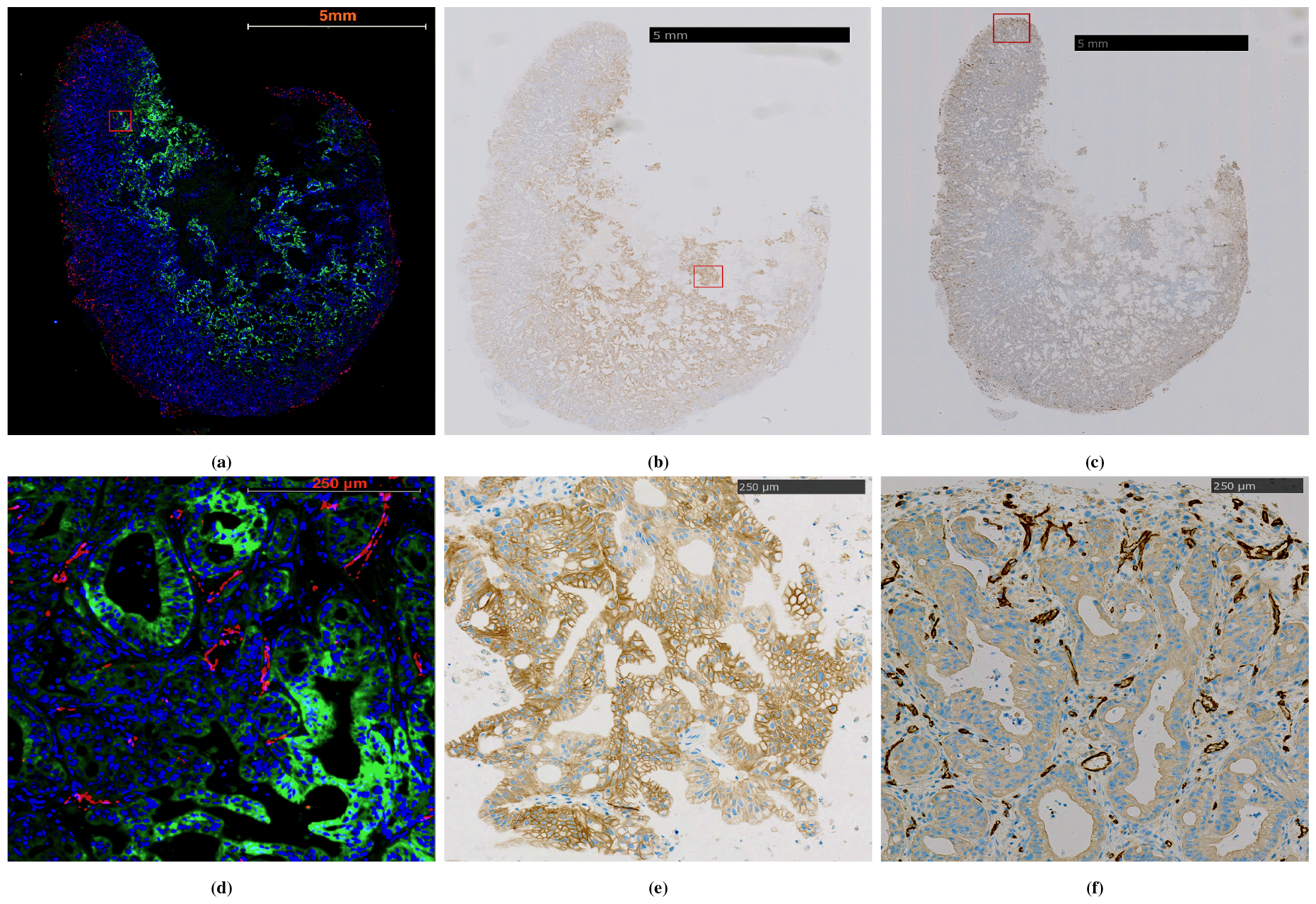
Our generated dataset comprises of images of tumor tissues stained with colors representing specific markers. Here we present the imaging data of pancreatic tumor tissue (1087), obtained through two distinct staining methods: immunofluorescence (IF) in the left column and immunohistochemistry (IHC) in the middle and right column images. The IF scanned image is depicted in fig. 1(a), and a zoomed-in section (highlighted by the red rectangle) of this image is shown in fig. 1(d). These images illustrate all the staining involved in our study, namely, CD31 in red, CA9 in green, and DAPI in blue, all on the same tissue. Figure 1(b), with a section zoomed in fig. 1(e), focuses explicitly on CA9 staining in brown, with color intensity indicating the gradient of hypoxia. Image 1(c) and its zoomed section in fig. 1(f) presents the blood vessels on the tissue adjacent to that used for CA9, also in brown. The cell nuclei are stained in blue color in the IHC-stained images, similar to the IF staining, as observed in the middle and right columns. It is important to note that although the tumor sample remains the same, the tissues involved in both scanning types differ.

### 2.2 Heterogeneity in data: Blood vessel diversity

Inter- and intra-tumoral heterogeneity is considered a challenging factor affecting therapeutic responses, and such heterogeneity is evident in our dataset, particularly concerning blood vessels, as depicted in fig. 2. The images reveal diversity in blood vessel architecture and geometry, showing long and large capillaries in fig. 2(a) and smaller ones in fig. 2(g). Capillaries are densely packed in patches fig. 2(g) and fig. 2(i), while they are rare in the second patch of fig. 2(d) and fig. 2(h). This diversity may arise from the underlying heterogeneity within and between tumors and the angle at which the 3D tissue is sliced for 2D imaging. Since effective oxygen distribution relies on blood vessel distribution in the tissue, quantifying this heterogeneity is crucial, especially in our modeling process discussed in the subsequent sections.

**Figure 2:**
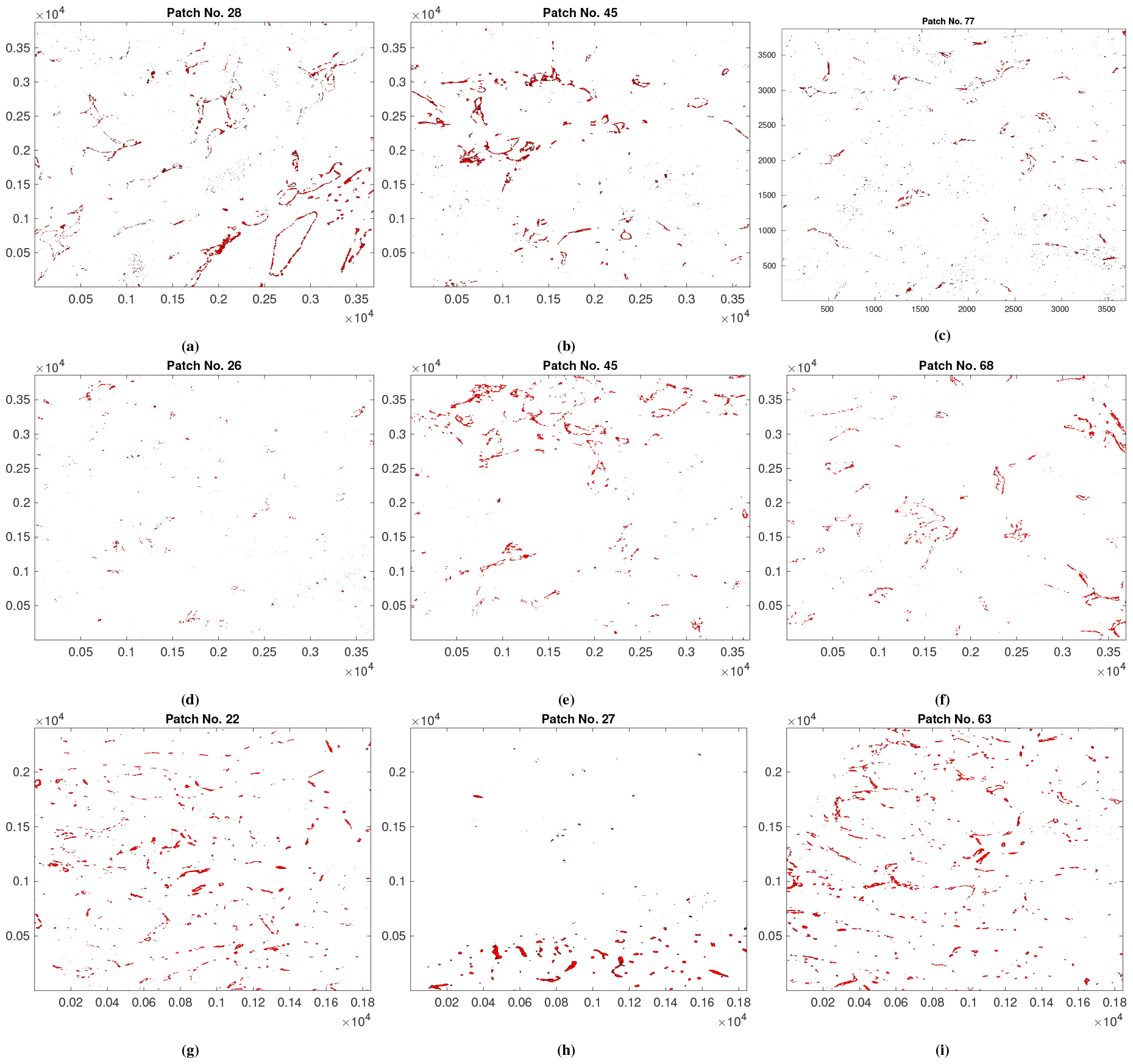
Various types of blood vessel architecture, represented by CD31 staining, are depicted in the top and middle rows for breast tumors (0520-04-1 and 0520-05-1) and in the bottom row for a pancreatic tumor (1087). These are patches of the mentioned tumor samples and are IF-type scans. For better visualization, the background of the IF images has been changed to white.

### 2.3 Data preprocessing

In order to utilize this data for spatial modeling, we apply some preprocessing techniques. Since the IHC stains of CA9 and CD31 are not conducted on the same samples, we perform image registration to obtain aligned images. Image registration is performed using two methods:

- Control point-based method: In this feature-based registration method, we manually identify a few matching features in both images (CD31 and CA9 stained IHC images) – specifically, matching tissue shapes. We choose 10 such points identical for each IHC scan pair. This selection of points is followed by estimating an affine transformation that differentiates one image (CD31 stained image, considered as the fixed one) from the other image (CA9 stained, considered as the moving one) geometrically. Applying the estimated transformation to the moving image aligns it with the fixed image.
- Optimization-based method: In this intensity-based registration method, we estimate the transformation using a regular step gradient descent optimization with mean square error metric configuration using the function `imregtform` in MATLAB^3^.

For the images utilized in this study, the control point-based method outperforms the intensity-based method. However, the latter proves useful when marking identical features in the images is challenging.

Following the completion of registration, we employ color thresholding in MATLAB to isolate specific marker colors in the tissue images. This allows us to obtain distinct stains for each sample; for samples stained with the IF method, color selection, and corresponding thresholding are executed in Zen software^4^ to achieve the same outcome. Furthermore, given that these images are high-pixel images (i.e., a single image can be 20k *×* 20k in height and width), we extract small, representative patches from the whole image, particularly focusing on areas of interest for the subsequent steps outlined in the following sections.

### 2.4 Extraction of marker densities

After the necessary processing, we obtain images for the respective markers for each tissue sample. The upper row of fig. 3 depicts the CD31, CA9, and DAPI marker staining images obtained from the preprocessing of the IF-based method for a patch of a breast tumor sample (1090). To facilitate modeling based on these images, spatial marker densities are required. This goal is achieved through three methods:

**Figure 3:**
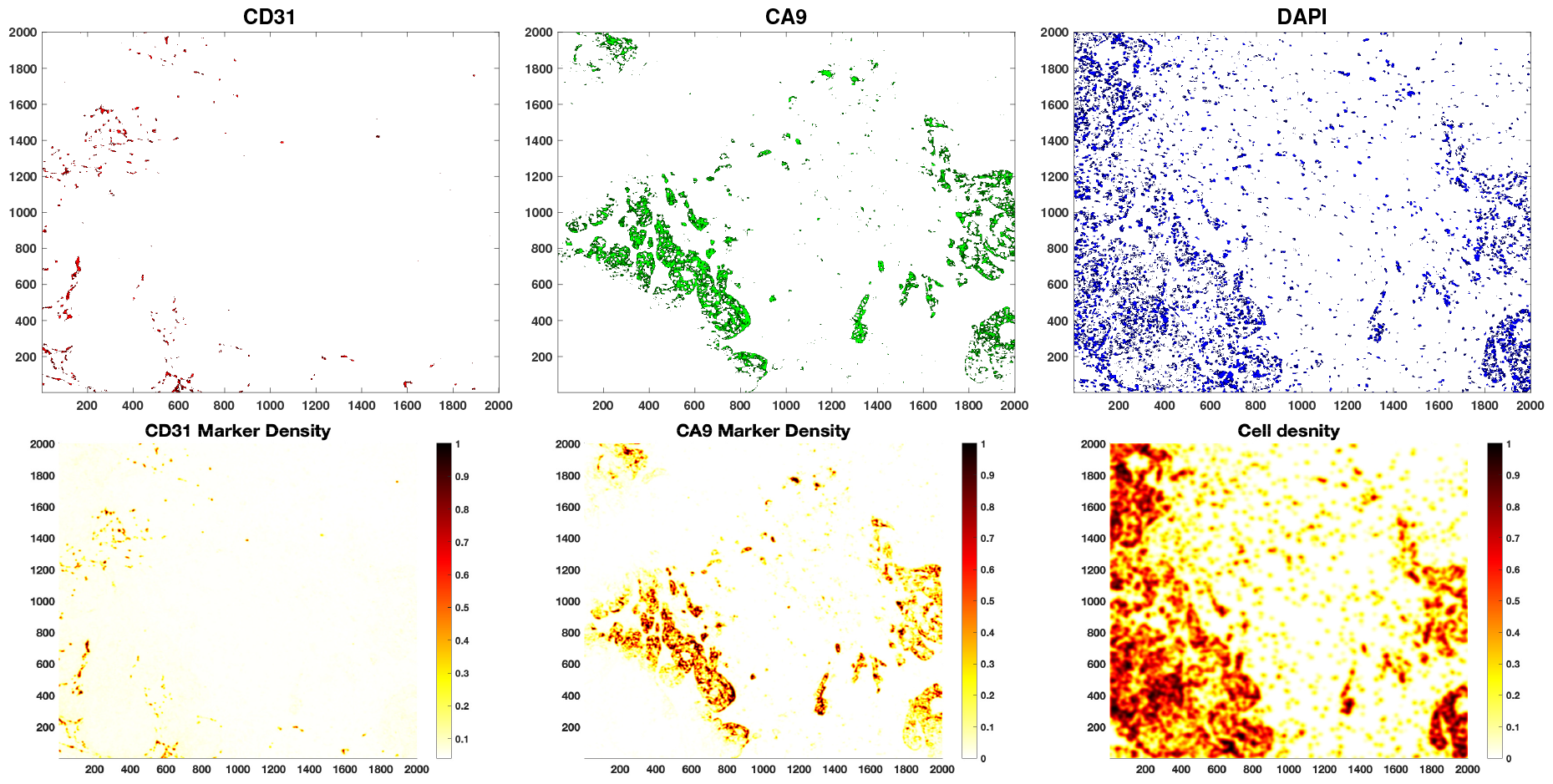
Upper row: Preprocessed image of a breast cancer tissue (1090) patch stained with CD31, CA9, and DAPI. Lower row: Corresponding densities.

- Intensity-based approach: In the first method, a 2D Gaussian filter^5^ (*G*) is applied to the images, resulting in pixel intensity-based density.
- Binary-pixel approach: The second method extends the first by obtaining a binary image with non-zero pixels for staining. Then, a Gaussian filter is applied to get the density.
- Kernel density estimator approach: The third method is based on kernel density estimate^6^ (KDE) for the image parts covered by the non-zero pixels. In this case, a Gaussian kernel is defined for each point^7^ on a considered grid in non-zero pixel regions, and the resulting density is the sum of all the kernels for the image.

The intensity-based approach is applied to extract density for CA9, as the intensity and gradient are meaningful to mark the level of hypoxia. The binary-pixel approach is used to obtain CD31 marker density. Initially, it is crucial to binarize the vascular and non-vascular regions for the blood vessels, followed by obtaining the density. Nuclei density, represented by the DAPI marker, is obtained using an area-based kernel technique. The resultant densities from both these methods are demonstrated in the lower row of fig. 3. The first two methods smoothen the images and make the density continuous, representing the continuous density of the blood vessels and hypoxia. Meanwhile, the third method captures the image portions covered by non-zero pixels and the proximity of the cell nuclei, forming a map of cells close to high density and low density for rarely packed cells.

### 2.5 Heterogeneity Quantification

To quantify the heterogeneity of blood vessels, we employ Claramunt entropy [8] within patches. This entropy metric, a modified version of the Shannon entropy, takes into account the spatial distribution of blood vessels. Assuming that an increase in distance between entities of the same type and a decrease in distance between entities of different types will result in increased entropy, the spatial entropy *H*_*s*_ is defined as follows:

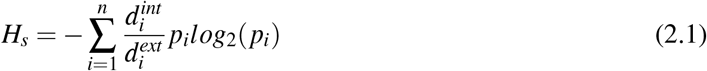

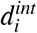 denotes the average distance between the entities (blood vessels) of the same type *i* (*i*.*e. i* intra-distance), while 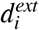 represents the average distance between the entities of a given type *i* and the entities of the all other types (*i*.*e*. extra-distance), here *p*_*i*_ denotes the proportion of each entity *i*, with *n* being the number of such entities.

To apply the entropy method previously mentioned, we first categorize image patches based on the geometrical variations of blood vessels present within them. These variations are determined by the number of pixels in a region covered by each capillary in a given patch. To ensure consistency across all images, we classify these variations into five categories (𝒞_*i*_) based on pixel area^8^ (𝒜_*i*_), *i* ∈ [1, 5] of blood vessels, ranging from very small to massive vessels.

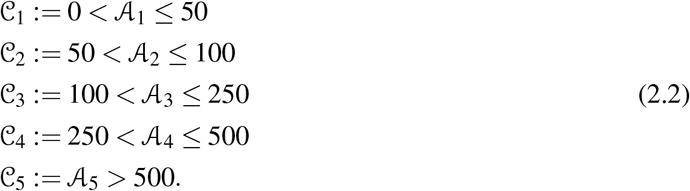

In the context of the formulation presented in eq. (2.1), the term *p*_*i*_ denotes the percentage of all non-zero pixels associated with the category 𝒞_*i*_. For a particular patch, as depicted in fig. 4(a), these categories are graphically represented using five distinct colors, as shown in fig. 4(b). Each color signifies a category 𝒞_*i*_, with the boundaries of individual blood capillaries drawn by these colors. Consequently, intra-distances 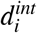 refer to the distances between vessels of the same color, whereas inter-distances 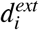 are the distances between vessels of different colors. The measurement of distances is achieved using the centroid of each vessel, identified as the connected pixels within the image. This approach comprehensively characterizes the spatial relationships between blood vessels, forming the basis for the subsequent entropy calculation outlined in eq. (2.1).

**Figure 4:**
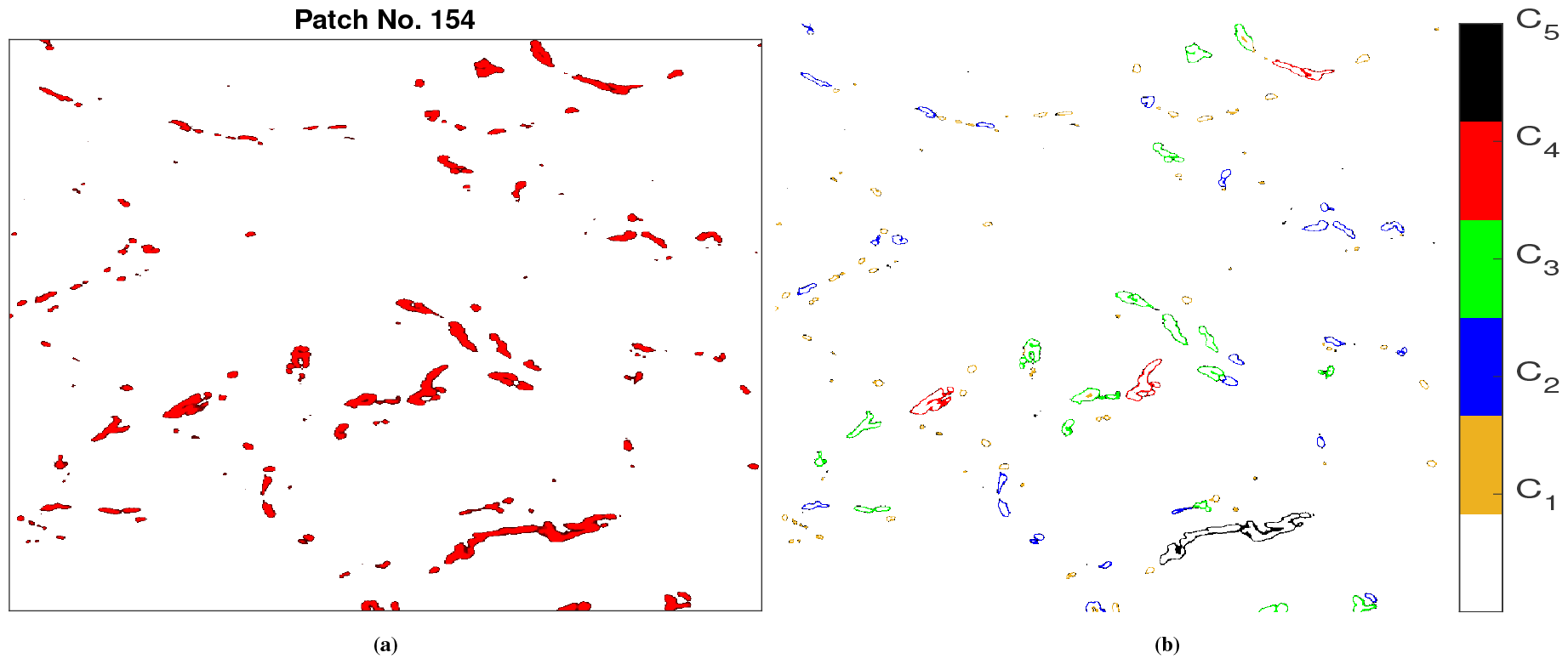
(a): Blood vessels of a patch (pancreatic tumor 1087), (b): Categorization based on pixel area covered by a blood vessel, recognized as a connected component; the colors correspond to the categories as represented by the color bar.

Applying a moving window approach, we compute the entropy for patches within each sample using eq. (2.1). As illustrated in fig. 5(a), we initially divide an image sample into 100 patches. Subsequently, we calculate the diversity index (*H*_*s*_) for each patch, as one of the patches exemplified in the zoomed-in form in fig. 5(b). The resulting diversity map for this tissue sample is depicted in fig. 5(c). The colormap in fig. 5(c) visually represents the diversity of blood vessel architecture within a tumor sample. This quantification of entropy provides valuable insights into the spatial distribution and variation of blood vessels, serving as a basis for identifying similar patches during the validation process.

**Figure 5:**
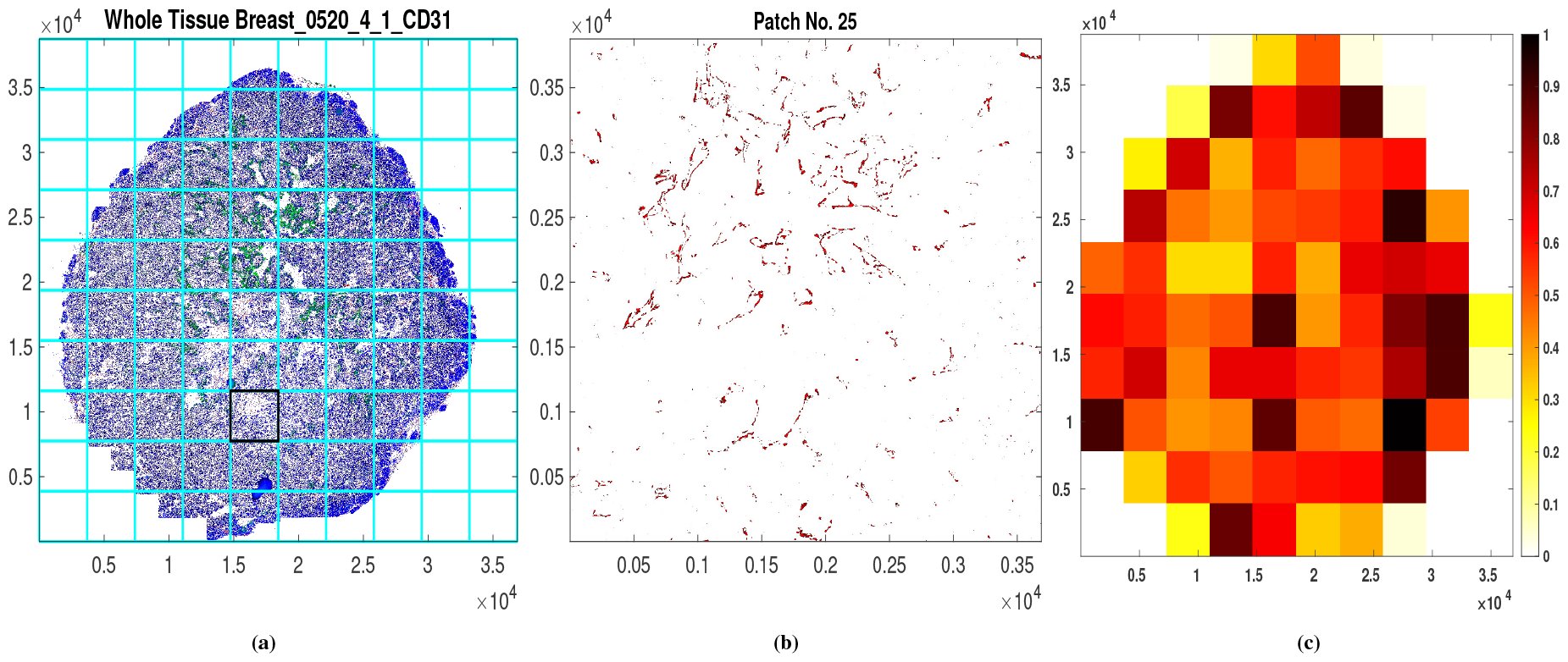
(a): A full sample of breast tumor blood vessel distribution, (b): a zoomed patch, (c): calculated heterogeneity score obtained using eq. (2.1). The patch numbering is executed in a Cartesian coordinate system manner, commencing from the bottom left and then progressing from left to right in the x-direction. This counting pattern repeats until reaching the top right corner.

### 2.6 Modeling

Utilizing the data mentioned above, which includes the densities of blood vessels, hypoxia, and cell nuclei, we develop a mathematical framework and physical models to investigate the relationship between oxygen and hypoxia distribution within tumor tissues. We begin by forming a model of oxygen distribution.

#### 2.6.1 Model of Oxygen Distribution

The dynamics of oxygen distribution within the microenvironment are influenced by various factors, including blood vessel architecture, geometry, functional state, permeability, the metabolic activity of cells, and oxygen consumption. In this study, we specifically focus on the following aspects:

- Production of *O*_2_: Our model assumes a direct correlation between oxygen production and blood vessel density in the tissue. Increased vessel density, particularly of capillaries, enhances oxygen exchange between blood and tissues. As detailed in section 2.4, estimating vessel density involves considering both overall density and spatial distribution. By incorporating vessel positions, we account for each vessel’s neighborhood, crucial for determining local oxygen supply.
- Neighborhood Determination: The appropriate neighborhood for *O*_2_ distribution is determined by the ability of *O*_2_ molecules to migrate the longest distance from their source vessels. To model this, we consider the random motion of oxygen molecules in the tissue, hence employing a linear diffusion to quantify this process. This method is a PDE counterpart of the experimentally observed diffusion length, which is applied in some previous studies as in [50] and [52].
- Consumption in the Tissue: Oxygen undergoes consumption in the tissue, primarily by cells and other chemical reactions in the tumor microenvironment. The uptake of oxygen is a crucial factor affected by the cellular metabolic state and the state of the tumor microenvironment. From a treatment perspective, a decrease in consumption can be utilized to reduce hypoxia [13]. In our modeling approach, we consider uptake directly proportional to the available oxygen concentration, encapsulating all the influencing factors into a unified representation.

To facilitate our modeling process, we introduce some notations for the variables involved. We consider the following:

- *u*(*t*, **x**): the oxygen concentration depending upon time *t* and space **x**,
- *D*: diffusion coefficient of *O*_2_ (a constant),
- *V* (**x**): blood vessel density,
- *α*: oxygen uptake coefficient,
- *β* : oxygen production coefficient.

Given the considerations outlined for modeling, along with the declared variables, we formulate a reaction-diffusion model for oxygen distribution in the following form:

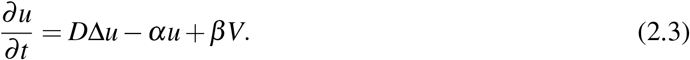

In this formulation, the modulation of oxygen concentration over time is governed by constant diffusion (represented by the first summand on the right-hand side of eq. (2.3)). Additionally, there is a constant uptake proportional to the existing *O*_2_ distribution (second summand) and oxygen production linked to the blood vessel density (third summand). This formulation aligns with previous models developed in [27, 54, 45]. Notably, in our data-driven approach, we have established an explicit relationship between blood vessel density obtained from data and oxygen production.

Further, as oxygen dynamics is rapid enough and it reaches the corresponding equilibrium state very quickly, as in [54, 27], we consider the steady state from the reaction-diffusion model:

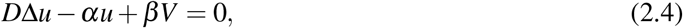

with boundary conditions: ∇*u n* = 0, which ensures no oxygen flux exchange between the considered domain and outside [54].

The model is eq. (2.3)) ensures the supply of oxygen by the available blood vessels; however, the production should not be unbounded. To incorporate this phenomenon, we define a limited production in another form of the model:

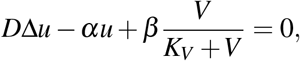

where *K*_*V*_ is the carrying capacity of the blood vessel density. Rewriting 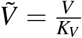 and dropping the tilde for simplicity, we get the steady-state form as:

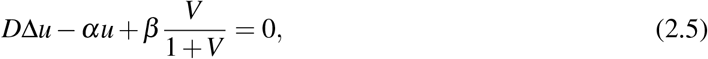

with the same no-flux boundary condition.

The uptake of *O*_2_ in eq. (2.5) is modeled as proportional to its availability in the tumor microenvironment. A linear relation between cell density and oxygen uptake has been observed in [25]. This uptake can be associated with the cell density (irrespective of being normal and tumor cells), with the following modified model

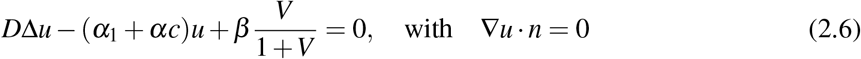

where *c*(**x**) denotes the cell density obtained from the DAPI marker staining, as explained in section 2.4 and *α*_1_ is a non-negative constant. However, the metabolic state of cells also plays an important factor in the consumption of *O*_2_.

#### 2.6.2 Modeling of Hypoxia-Oxygen Relationship

As mentioned in the introduction, hypoxic regions result from poor oxygenation. Therefore, the relationship between hypoxia and *O*_2_ would be formulated in an inversely proportional manner.

One initial choice for modeling this relation is to consider a linear decay of hypoxia with an increase in *O*_2_. Let *h*(*u*) represent hypoxia as a function of normalized oxygen distribution, and the linear decay can be expressed as:

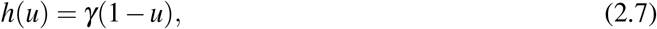

here, *γ* is a dimensionless constant conversion coefficient.

Another formulation could involve an exponential decay:

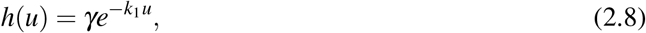

with *k*_1_ being a dimensionless constant representing the rate of decay. Such decay has also been observed in [40].

A more general approach is to consider a sigmoidal form:

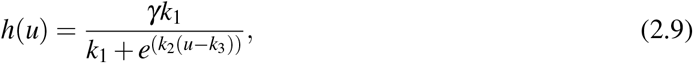

In this case, *k*_*i*_, where *i ∈*[1, 3], are constants that govern a more controlled decrement of oxygen leading to hypoxia.

Utilizing the three models mentioned above, we obtain the corresponding hypoxia profiles, each representing the decay of hypoxia (linear, exponential, or sigmoidal). Furthermore, a switch-like^9^ relationship between *O*_2_ and hypoxia is proposed in previous works, such as in references [7, 37, 14]. We incorporate this response with the lower and upper thresholds of *O*_2_ corresponding to non-zero hypoxia, denoted as *O*_*l*_ and *O*_*h*_; this relationship is modeled as follows:

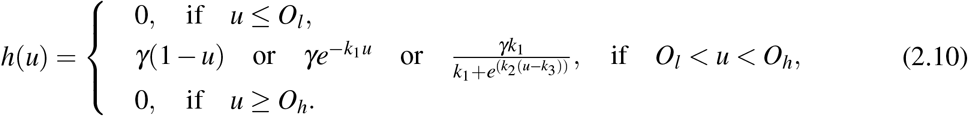

For the particular values of *O*_2_ thresholds (*O*_*l*_ = 0.2 and *O*_*h*_ = 0.8, fig. 6 illustrates the hypoxia-*O*_2_ relationship represented by (2.10). The upper bound (*O*_*h*_) is evident as hypoxia decreases to zero in the presence of sufficient oxygen (normalized value = 1). This decrease can manifest as a linear decay (fig. 6(a)), a sharp exponential decay (fig. 6(b)), or a sigmoidal-type decay (fig. 6(c)). The lower bound (*O*_*l*_) models two processes. First, in cases of severe and prolonged hypoxia, HIF-1 levels decline, as suggested in [14, 29, 41]. As HIF regulates CA9 [20, 33], which is a well-characterized target of HIF-1 [5]. Hence, below the lower threshold (*O*_*l*_), the hypoxia *h*(*u*) switches to zero. Secondly, this lower bound is also useful to incorporate prediction for non-cellular regions, where CA9 and CD31 markers are absent simultaneously. However, in our simulations, these parameters are obtained from the underlying data using an estimation method.

**Figure 6:**
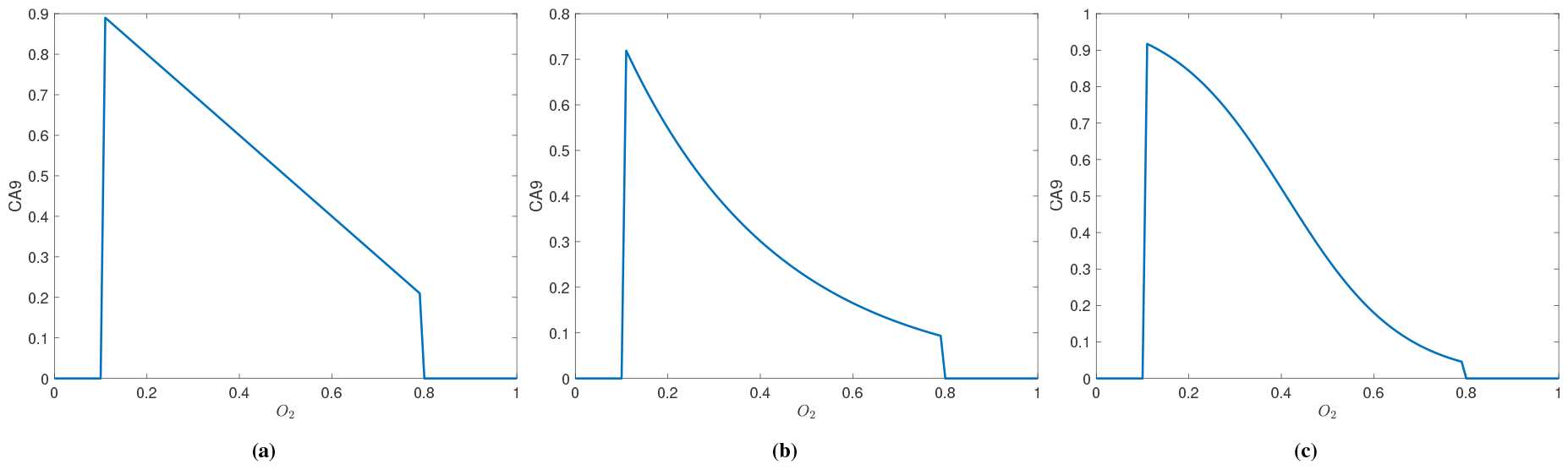
1D representation of hypoxia-*O*_2_ relationship formulated in (2.10).

#### 2.6.3 Final model system

Hence, our developed model system involves a PDE (eq. (2.4)) for oxygen distribution and an algebraic relation between *O*_2_ and hypoxia profile in (eq. (2.10)) based on the mentioned dataset, which will be utilized to estimate the involved parameters. This estimation is performed with the exponential decay form of equation eq. (2.10). We have also tested the model with bounded production described by (2.5) but did not observe an increase in training and validation performances.

### 2.7 Numerical Simulation Methods

To estimate the parameters in our formulated system, we numerically solve the PDE presented in eq. (2.3). We use a finite-difference scheme where the spatial domain is discretized into 400 *×*400 grid points. A five-point stencil (central-difference) discretization is applied to the diffusion term on these grid points. The resultant system of linear equations consists of 160,000 unknowns, representing the values of unknown (*u*(**x**)) at the grid points. The simulations are conducted within the MATLAB environment.

Upon solving the PDE in eq. (2.3) for a given set of parameters, the *O*_2_ distribution *u* is obtained at the discretized points. Subsequently, hypoxia *h*(*u*) is calculated using eq. (2.10) at the considered grid points within the domain. This forms the basis for further analysis and parameter estimation.

### 2.8 Estimation of parameters

Model parameters are determined using an optimization method. To do this, we first define a loss function as:

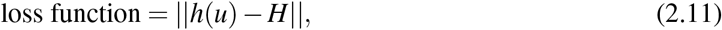

where *h*(*u*) is the computed hypoxia from (2.10), *H* represents the hypoxia profile obtained from the CA9 scan and | |. | |denotes the square of *L*^2^-norm per grid point. The loss function is computed within the considered domain where either of the scans (CD31 or CA9) is non-zero. The minimization of this loss function provides estimates for the parameters.

The optimization is performed in MATLAB using the built-in `MultiSearch` function, where 100 to 500 random values are chosen for the initial set of parameters in the optimization process. This function uses multiple random starting points and generates optimal and also sub-optimal solutions. The search of optima is parallelized and calls upon the sequential quadratic programming (sqp) algorithm via `fmincon` in MATLAB. Additionally, we have also explored the non-linear least square fit method and `GlobalSearch` algorithm, which is based on the scatter-search method of generating trial points and automatically selecting 1000 starting points. Our findings indicate that the `MultiSearch` with `fmincon` equipped with the sqp algorithm outperforms other methods, at least as far as our problem is concerned.

### 2.9 Training and Validation Datasets

We divide the dataset into two sections: one for training the parameters and the other for validating the obtained model predictions. In this partition, we ensure diversity in both datasets by including all types of tumor samples and scan types. As illustrated in the table in table 2, we train on the IF samples for convenience, as they involve the same tissue for staining. Additionally, for validation, as depicted in table 3, we include both types of scans and tumor types to test the predictive capabilities of our modeling framework thoroughly. Next, we further divide each sample into small patches and train the model on each of them, as shown in table 2, indicating the number of patches.

**Table 2:** Training Dataset.

| Sample Name | Tumor Type | Tissue | # sections | Method | No. of Patches |
| --- | --- | --- | --- | --- | --- |
| 1092 | Breast | PDX | 1 | IF | 81 |
| 0056 | Breast | PDX | 1 | IF | 76 |
| 0520 | Breast | PDX | 53 | IF | 4045 |
| 1137 | Pancreatic | PDX | 1 | IF | 77 |
| 1905 | Pancreatic | PDX | 1 | IF | 72 |

**Table 3:** Validation Dataset.

| Sample Name | Tumor Type | Tissue | # sections | Method |
| --- | --- | --- | --- | --- |
| 1090 | Breast | PDX | 1 | IF |
| 1092 | Breast | PDX | 1 | IHC |
| 1087 | Pancreatic | PDX | 1 | IF, IHC |
| 1905 | Pancreatic | PDX | 1 | IHC |
| 0721 | Ovarian | PDX | 1 | IHC |

### 2.10 Training

As depicted in fig. 5(a), we partition each image into 100 patches and independently train the parameters for each patch. To compare against a worst-case scenario, we evaluate the minimized loss function against the mean of 50 random values of the hypoxia profile at each grid point. Further, in the training process, we record the parameter values corresponding to the 50% higher than the global optimum for each patch. Executing this process for the entire training set enables us to obtain optimal and sub-optimal parameters and an interval for each parameter for each patch, considering sub-optimal solutions 50% higher than the global optimum. This information provides insights into the uncertainty of the estimated parameters across different patches in the image.

### 2.11 Stratification of tissue regions

Upon completing the training phase, the trained framework is applied to classify a patch and identify similar patches from the trained dataset based on the blood vessel architecture. For a given patch from a tissue sample outside the training data, the classification process involves the following steps:

- Assigning Heterogeneity Score (*H*_*s*_): Initially, calculate the heterogeneity score for the patch’s blood vessel density using the method outlined in section 2.5 to quantify hypoxic regions.
- Proportion of Non-zero Pixel (*N*_*p*_) Calculation: Next, we calculate the proportion of non-zero pixels representing blood vessels in the patch. As discussed in section 2.4, a binary pixel method is used to detect blood vessels in the images, providing a measure of the vessel-filled region in the patch.
- Initial Interval Determination: Determine two intervals with a variance of ±10% above and below both calculated values: (*max*(0, *N*_*p*_ *−* 0.1*N*_*p*_), *N*_*p*_ *±*+ 0.1*N*_*p*_) and (*max*(0, − *H*_*s*_ 0.1*H*_*s*_), *H*_*s*_ + 0.1*H*_*s*_), ensuring that the minimum values remain non-negative.
- Identification of similar patches: Once heterogeneity and non-zero pixels intervals are obtained for the target patch, identify patches with trained parameters, whose two scores lie within the intervals. These identified patches are considered “similar” patches, and their parameters can be utilized to predict the hypoxia profile. Initiate an iterative search starting with the initial interval. The algorithm searches for trained patches within this interval. The search iteratively increases the interval until there is at least one set of values satisfying both conditions specified by these two intervals, providing a similar patch.

Hence, this classification ensures that similar patches contain equivalent vascular regions and that their spatial spread is comparable.

### 2.12 Validation of the physical model

This step involves predicting the hypoxia profile for a patch and subsequently for an entire image, then comparing the predictions with the actual data. The process is as follows:

#### 2.12.1 Validation for a patch

To predict the hypoxia profile of a patch, the following steps are undertaken, starting from image preprocessing to solving a PDE, ultimately providing insights into the hypoxia profile of the tumor tissue sample.

- Input Image: Tumor tissue image stained with CD31 and DAPI, representing blood vessels and cell nuclei, respectively.
- Necessary Preprocessing: Implementation of necessary preprocessing steps as mentioned in section 2.1 to obtain the respective markers’ densities.
- Patch Extraction: We divide the input sample into small patches. Depending upon the image pixel size, we divide them into 25 to 100 small patches.
- Classification of a Patch: Next, we apply the stratification method to identify similar patches from the training dataset, thereby obtaining an interval for each parameter.
- Parameter Assignment for Each Patch: We obtain a range of parameters for the patch, and each value can be identified as a different snapshot of dynamic oxygenation. For a general case, parameters are assigned for the patch by taking the mean of the interval of the similarly trained patches.
- PDE Solution: With the obtained parameters, the PDE in eq. (2.4) is numerically simulated for the patch to get the *O*_2_ distribution.
- Cell Region Identification: Identification of the cell regions based on DAPI marker density is performed to mark the regions where cells can be present and to make predictions for hypoxia.
- Hypoxia Calculation: Application of the hypoxia equation eq. (2.10) and then setting the obtained to zero for non-cellular regions provide the desired profile for hypoxia for the input patch.
- Result Image: The final result is the hypoxia profile for the input patch.

#### 2.12.2 Validation on whole sample

We assume constant parameters in the formulated model system described by eq. (2.3) and eq. (2.10), which may not hold for all tumor tissue samples. We adopt a data-driven approach to retrieve space-dependent model parameters from the data to predict the hypoxia profile for an entire sample. This is accomplished through the following steps using a moving window approach:

- Partition the input sample into several patches (e.g., 100 patches per sample), as illustrated in fig. 5(a).
- Apply a classification strategy based on the heterogeneity score of each patch, as detailed in section 2.2.
- For each patch, acquire parameters from similar patches in trained samples, following the process detailed for the single patch case.
- The obtained parameters for each patch result in space-dependent parameters that remain invariant within a patch, expressed by the equation:

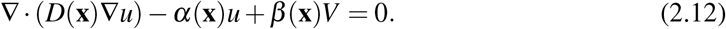
- Together with the no-flux boundary condition, eq. (2.12) serves as a data-driven, space-dependent counterpart of eq. (2.4).

This adaptation enables the consideration of spatial heterogeneity in parameters within the tissue sample, providing a more realistic representation of oxygen and hypoxia profiles. To address the spatial dependence of the diffusion coefficient *D* and prevent numerical oscillations during the simulations of eq. (2.12), we employ a conservative form of the five-stencil discretization method, as described in section 2.7.

## 3 Results

### 3.1 Training the mechanistic model for oxygen-hypoxia relationship

A simple yet effective mathematical framework is formulated to predict hypoxia from a known vessel architecture. The framework is mechanistic and data-oriented, utilizing underlying data for model development, training, and subsequent validation. The approach combines a theoretical foundation with empirical insights from the available dataset, emphasizing a balance between mechanistic understanding and practical applicability.

Following the training protocol detailed in section 2.10, it is apparent that the proposed framework aligns effectively with the data, as evidenced by the representations in Figures 7 and S2. These scan images result from two distinct staining techniques, as elucidated in section 2.1, and the modeling framework exhibits a robust fit with both types of scans. Beyond the visual appeal of the fit comparison, a quantitative assessment reveals a strong agreement as we compare the minimized loss function values with the worst-case scenario, as explained in section 2.10. This is observable in the figures presented in 7 and S2. This consistent trend persists across the entire training set, where each patch’s minimized loss function value is significantly smaller than the worst values, as shown in the fig. S1.

**Figure 7:**
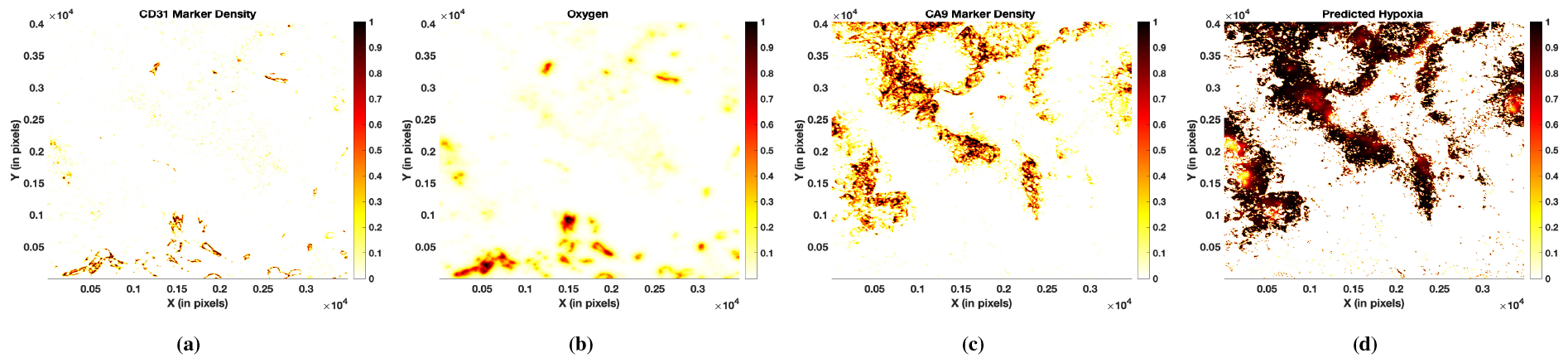
Data fitting and parameter estimation for a patch extracted from the breast tumor (0520-4-2) tissue image, an IF scan, involve utilizing the training procedure outlined in section 2.10 and the specified optimization method for extracting parameters. Initially, we determine the blood vessel density from the CD31 scan, shown in (a), and the hypoxia profile from the CA9 marker in (d), following the details in section 2.4. Further, with the initial guess of parameters, we calculate the oxygen distribution using the obtained blood vessel density and apply eq. (2.3). The process continues with acquiring the hypoxia profile using eq. (2.10). The obtained hypoxia profile is then compared with the CA9 marker density in (c) through the loss function in eq. (2.11). Beginning with the initial parameter guess and iterating through this process until the loss function in eq. (2.11) is minimized, an optimal value is achieved. The resulting oxygen and hypoxia distribution, depicted in (b) and (d) respectively, with the loss function value of 0.07 and the worst case value being 10.59.

### 3.2 Biological heterogeneity and epistemic uncertainty are responsible for distributed physical parameters

Upon estimating parameters for all patches within the training set, a distribution for each parameter is obtained, as depicted in Figure 8(a). This visualization shows the parameter distribution, revealing notable inter-patch heterogeneity. The observed hypoxia heterogeneity in experiments [24] is consistent with the model predictions, particularly in terms of inter-patch variation of parameters.

**Figure 8:**
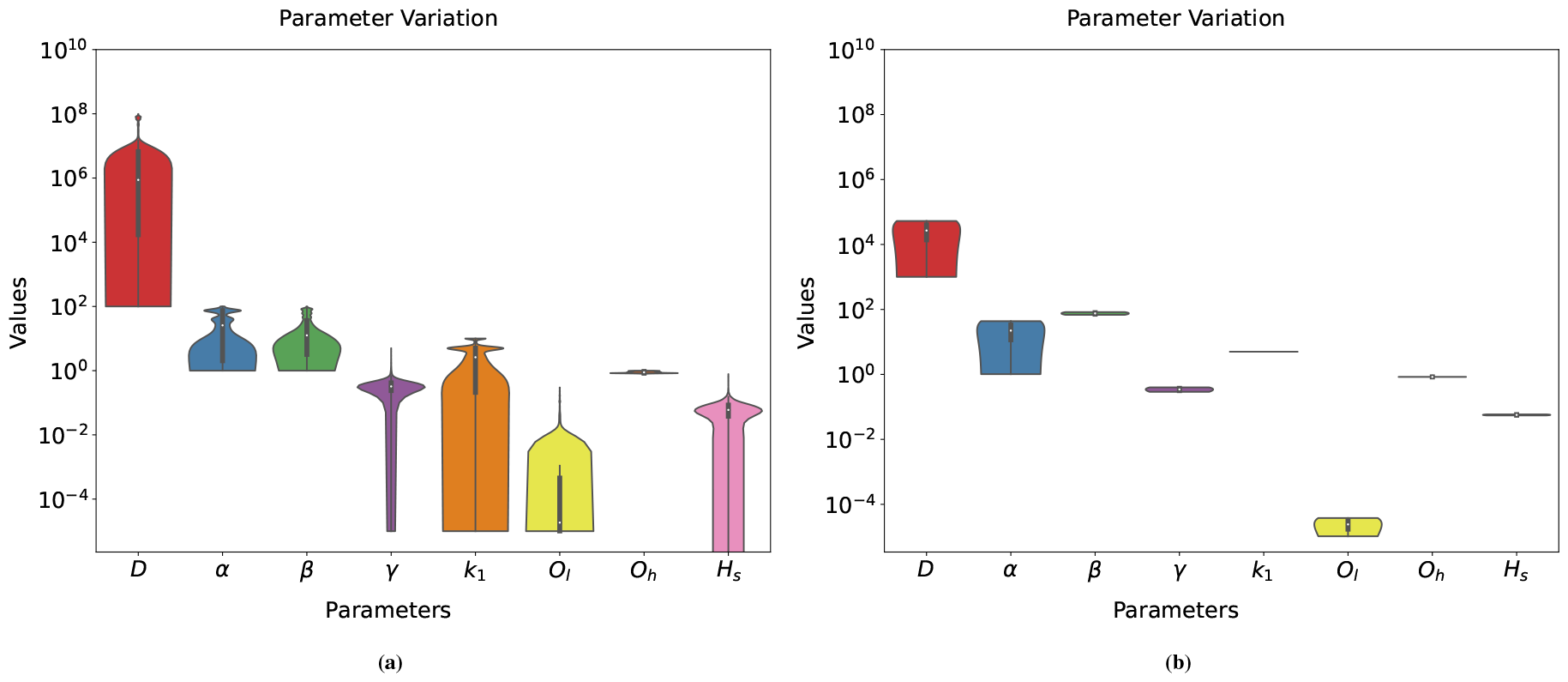
Inter-patch parameter variation and the reduction of this variation through stratification of patches. (a): The trained parameters exhibit variation, as shown in the violin plots from left to right: diffusion coefficient *D, O*_2_ uptake and production rates (*α, β*), *O*_*l*_, *O*_*h*_, and heterogeneity score for the trained patches. Notably, these distributions are often multimodal. The variability in *D* spans several orders of magnitude, showcasing highly and poorly diffusing *O*_2_ molecules. The multimodal nature of production and uptake rates reflects variations in cellular demands. Meanwhile, the modality in *k*_1_ represents the variation in the decay of hypoxia profiles between different patches. (b): Distribution of parameters obtained after the classification of a selected patch from a breast tumor (0520-25-2) scan of the IF type. The parameter intervals are determined from similar patches, leading to significantly reduced variability compared to the distribution shown in (a). The parameters include the diffusion coefficient *D, O*_2_ uptake and production rates (*α, β*), *O*_*l*_, *O*_*h*_, and heterogeneity scores for the trained samples, arranged from left to right.

The first source of parametric variability is biological and integrates several factors. Tissue oxygenation is a complex process influenced by various factors beyond our modeling constraints, with one such factor being the dynamic nature of oxygenation, particularly in vascularization induced by normal or tumor-related processes. This intricate process involves a series of coordinated steps, commencing with tumor cells overexpressing hypoxia-inducible regulators of angiogenesis, including vascular endothelial growth factor (VEGF). As a chemoattractant, VEGF promotes the proliferation of endothelial cells (ECs) lining the surrounding blood vessels. For a more detailed model and qualitative insights, particularly in the context of glioma, we refer to [31].

The complexity of this process and the diversity in blood vessels and hypoxia profile are mirrored in the parameters. The observed inter-patch or inter-tumor variability is intricately linked to the underlying dynamics, facilitating an exploration of different possibilities and leveraging the mechanistic behavior of the model. These obtained distributions provide insights into the diverse, dynamic states of tissue oxygenation, as elaborated in subsequent sections.

Another source of parametric variability is epistemic, stemming from the fact that several parameter sets can result in similar fits. Therefore, trained parameters are inherently uncertain. In the optimization process, parameters corresponding to the minimum loss function, as defined in eq. (2.11), are selected. However, introducing a bit more tolerance in the loss function results in an interval of parameters, reflecting epistemic parametric uncertainty. This type of variability results from insufficient training data, which can be mitigated by adding more data.

### 3.3 Prediction of hypoxia from stained tissue images using the physical model

#### 3.3.1 The stratification, based on blood vessel heterogeneity, reduces the variability of parameters

As we explore the variability of parameters across patches and consider the underlying dynamics, it becomes essential to narrow down options when making predictions using the framework. Indeed, model predictions depend on parameters that are a priori unknown. Our strategy is to use the heterogeneity score (spatial entropy *H*_*s*_), which can be directly extracted from stained tissue images, to reduce the range of parameters required for prediction.

To assess the feasibility of this strategy, we employ the stratification method outlined in section 2.11 to associate a set of trained parameters with each new image patch for which we require prediction of hypoxia. This method effectively restricts the parameter range during the validation process. Applying the parameter assignment method, as detailed in section 2.12, reveals reduced variability in the parameters for individual patches—for example, one from breast tumor tissue (0520-25-2) and another from pancreatic tumor tissue (1087)—in the validation set. The former tissue image is an IF scan, while the latter is an IHC scan image. This reduced variability observation is evident from the distribution plots shown in fig. 8(b) and fig. S3.

#### 3.3.2 Prediction based on a restricted set of parameters and assignment of a state of vascularization

Using the stratification method, we narrow down parameter intervals, resulting in a reduced range for each parameter. While not providing an exact value, this interval offers a range of possibilities reflecting the underlying dynamics in the tissue. The variations in individual parameters signify different underlying dynamics in the concerned tissue. To better understand these dynamics, we analyze the changes in specific parameters one at a time while keeping the other parameters at the mean of their obtained interval for a patch sample. For the given blood vessel in fig. 9(a) of the patch, oxygen and hypoxia distribution is predicted, as shown in Figures 9(b) and 9(d), whereas the CA9 marker density is depicted in fig. 9(c). By exploring the range of parameters obtained for this patch, we observe the following:

**Figure 9:**
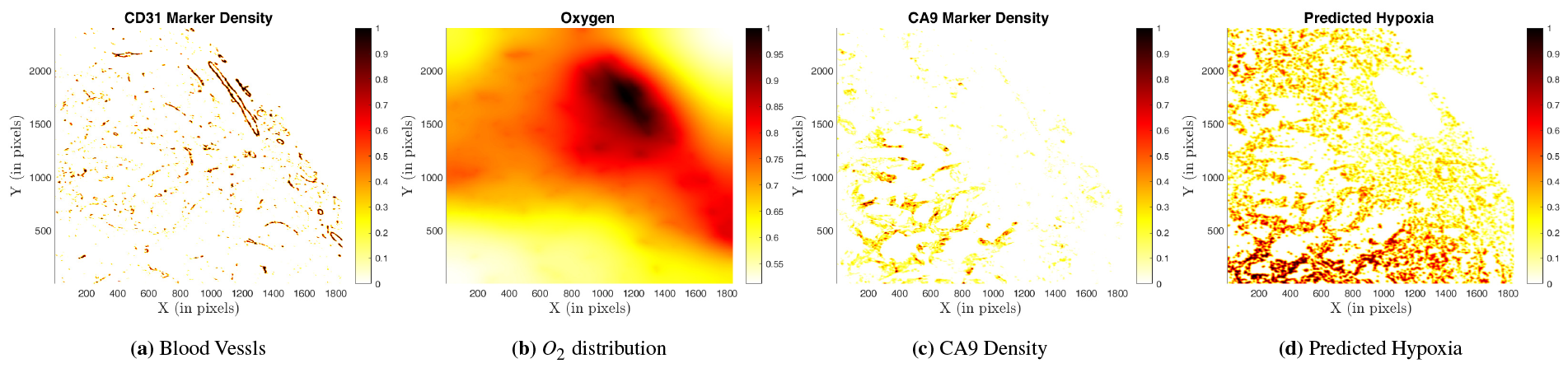
Prediction of oxygen distribution and the hypoxia profile for a patch of a pancreatic tumor sample (1087) of the IF type. The parameters used in this prediction are the mean values obtained from similar patches after the classification process.

- Diffusion coefficient (*D*): As mentioned, oxygen supply is assumed to be the diffusion of oxygen molecules produced by blood vessels. This diffusion is controlled by the diffusion coefficient (*D*). In a properly oxygenated tissue microenvironment, *O*_2_ molecules would be acquiring the maximum limit of the diffusion coefficient, resulting in less hypoxia, as seen in fig. 10(a). Conversely, less diffusion would lead to more hypoxia, as depicted in fig. 10(d).

**Figure 10:**
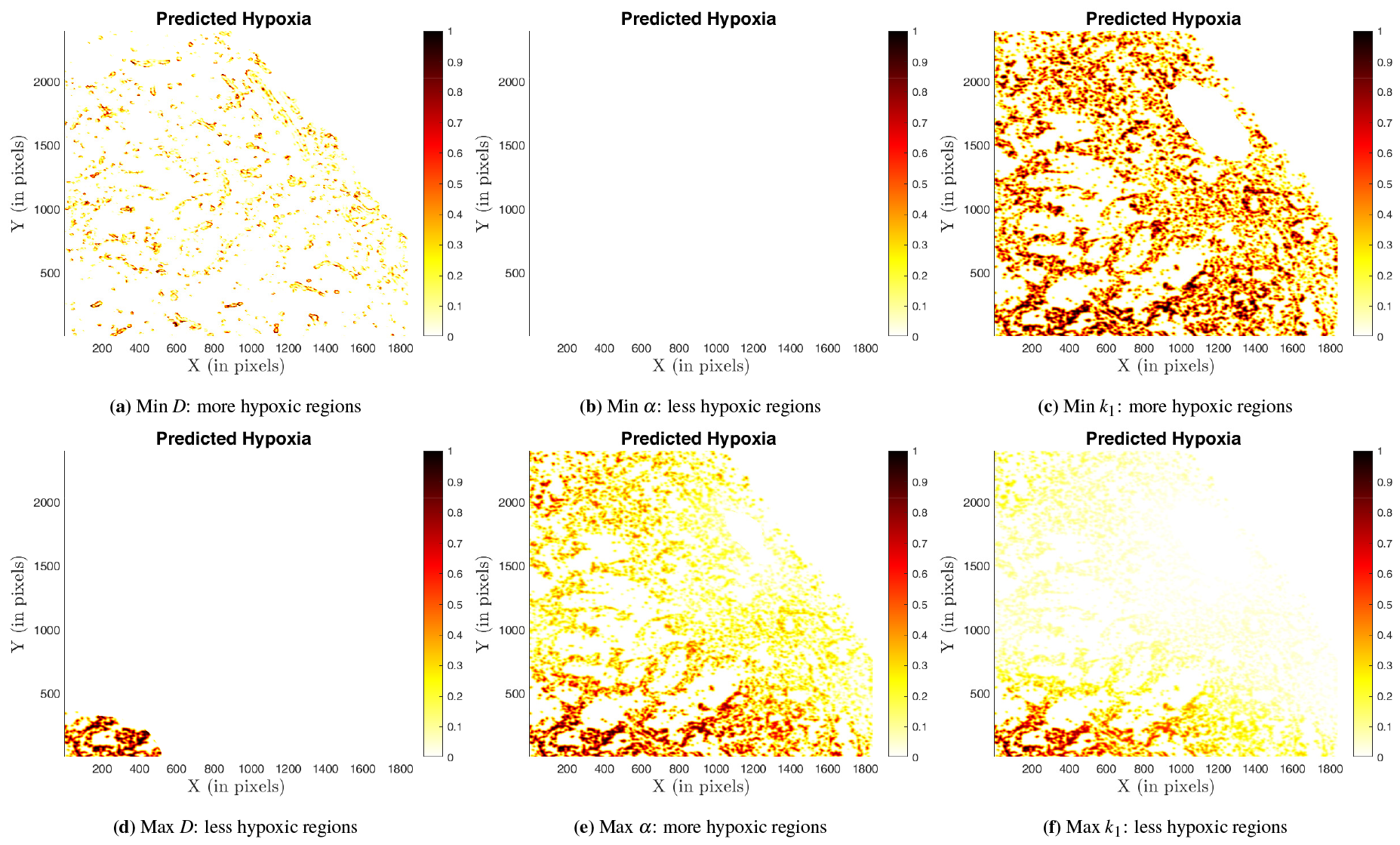
Impact of varying different parameters, each representing distinct dynamics within the intervals obtained after stratification and assigning intervals for each of them. Each subfigure individually displays the variation of a specific parameter while keeping the other values as the mean of the obtained ones after stratification. The results feature the same patch as shown in fig. 9. For example, a minimum diffusion coefficient results in more regions of hypoxia for the cells (fig. 10(a)), while very high diffusion would eliminate the possibility of hypoxic cells for most regions (fig. 10(d)). Similarly, deficient oxygen uptake would maintain the tissue normoxic (fig. 10(b)), whereas high uptake would result in hypoxic regions (fig. 10(e)). Additionally, an increase in the decay rate of hypoxia with oxygen would lead to hypoxic regions (fig. 10(c)), while a decrease shows the reverse trend (fig. 10(f)). Therefore, the variation in these classified coefficients for similar patches can be seen as a snapshot of different oxygen dynamics.
- Production and uptake coefficients: The production coefficient (*β*), related to blood vessels, characterizes the state of blood vessel functioning. Higher values of *β* reflect increased oxygen production, resulting in lower hypoxia. On the other hand, the Oxygen uptake (*α*) rate is modulated by the tumor microenvironment and cellular metabolism and plays a crucial role in the hypoxia profile. Increased uptake leads to more hypoxia, as demonstrated in fig. 10(b) for a minimum value and for a maximum value of *α* in fig. 10(e).
- Decay coefficient (*k*_1_) and conversion coefficient (*γ*): The decay coefficient *k*_1_, which signifies the rate of decay of CA9 marker with increasing oxygen concentration, exhibits a high hypoxic response at minimum values (fig. 10(f)), whereas maximum values result in low hypoxia (fig. 10(c)). A high value of *k*_1_, such as the maximum selected parameter, suggests a microenvironment where hypoxia diminishes rapidly with the concentration of oxygen. This could indicate a tissue state with strong oxygen needs, that reacts abruptly to a lack of oxygen. Conversely, a low value represents a scenario where the tissue is less dependent on oxygen, as in the case of anaerobic metabolism. As *γ* is a conversion coefficient limiting the maximum value of hypoxia distribution, a high value of *γ* implies a state with high oxygen demands, while a lower value suggests lower oxygen dependence.

Combining these parameters can help profile the blood vessel’s functioning, with several possibilities of parameter combinations reflecting different dynamic conditions of tumor oxygenation. Here, we focus on two scenarios: functional and non-functional blood vessels. Functional blood vessels may result from efficient diffusion, low oxygen uptake in tumor tissues, and a high decay coefficient of hypoxia with increasing *O*_2_. In contrast, the non-functional category includes less diffusion, less production, and high oxygen uptake—where even with high production, this could be the case for new vessel linings. The plots in fig. 10 showcase plots corresponding to these scenarios, extracting the dynamic nature of the dynamics through this static model.

### 3.4 Validation of the physical model using diverse tumor samples

A desirable feature of a mathematical modeling framework is its predictive nature, coupled with a mechanistic explanation. Moreover, it is crucial to validate these predictions to ensure biological relevance. As detailed in section 2.12, we validate the model predictions using the outlined procedure, beginning with the patch validation. In this section, we present the results, demonstrating alignment with the mean values of the obtained parameters. Although we have intervals for each parameter, the validation results are consistent with the mean values for many of these samples. The validation was performed using a diverse set of tumor samples (see Table 3).

#### 3.4.1 Validation for patches

Our model predictions demonstrate strong alignment with the validation data, as evident in figures figs. 11(a) to 11(d) and fig. S4 for IF, and figure fig. S6 for the IHC scan type. The predicted hypoxia profiles in fig. 11(d) and fig. S4(d) showcase the recovery of CA9 positive cells, utilizing the mean of the parameters after the classification process detailed in the previous section. While most CA9-positive cells are well-recovered, there are some additional tissue regions with cells marked as hypoxia-positive. However, these instances are relatively low compared to the well-aligned predictions. These additional positive markings may arise from various approximations and simplifications incorporated in our modeling process.

**Figure 11:**
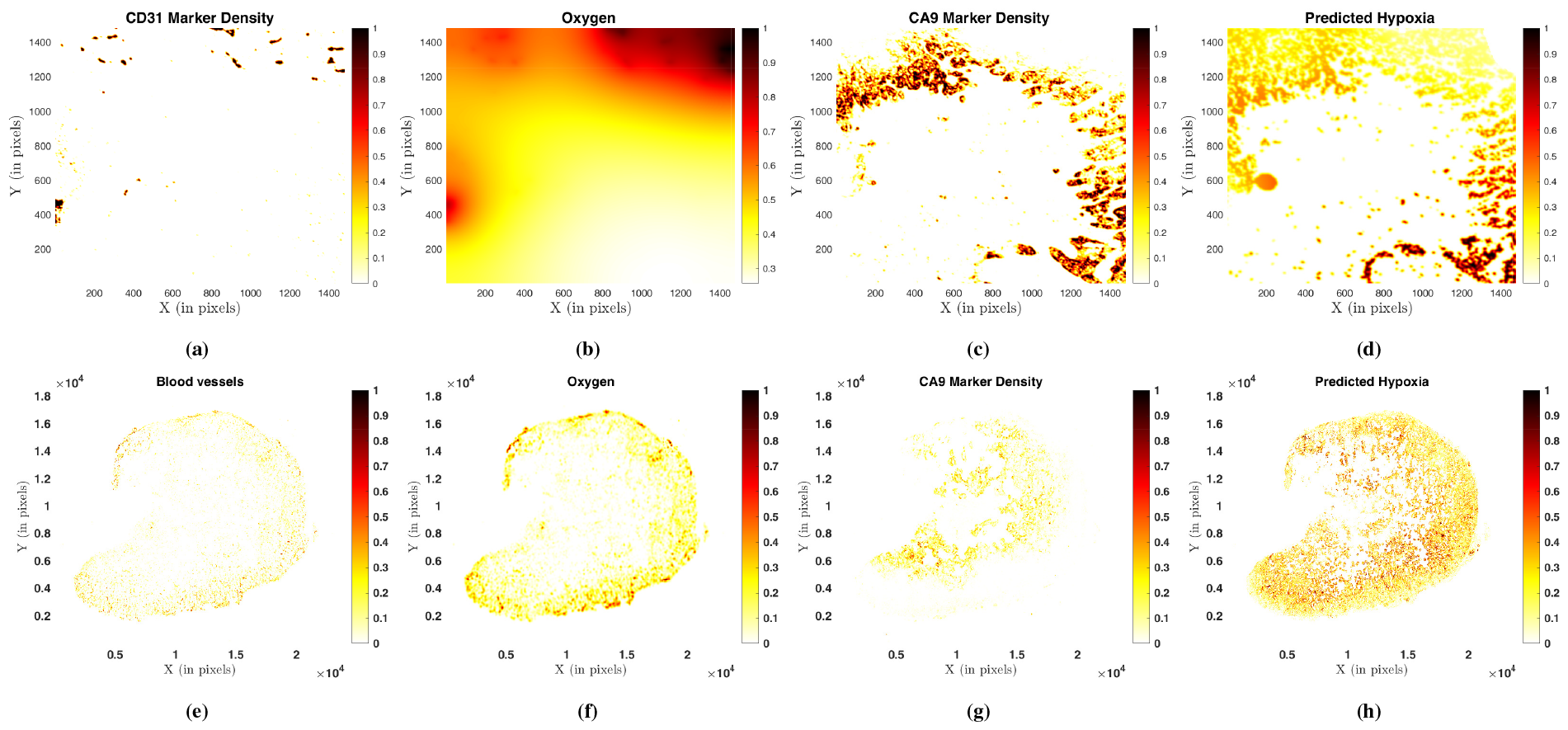
Upper row: Validation of the prediction for a patch of the breast tumor sample (1090), stained by IF, follows the steps outlined in 2.12. The blood vessel density is depicted in (a), and a range of involved coefficients are obtained. The resulting predictions from eq. (2.3) and (2.10), representing oxygen distribution and the hypoxia profile, are shown in plots (b) and (d), respectively. The hypoxia profile closely aligns with the CA9 density shown in plot (c), yielding a validation error 0.2310. **Lower row**: Predictions and their validation for the entire tissue sample were carried out following the steps detailed in section 2.12.2. Utilizing the blood vessel density from an input image, as shown in plot for a pancreatic tumor sample (1087) stained with IF, the model framework produces predictions for *O*_2_ and the hypoxia profile, presented in plots (f) and (h), respectively. Upon comparison with the CA9 data, displayed in (g), additional hypoxic cells are observed, resulting in a relatively high validation error of 0.6366.

Despite the presence of some additional cells marked as CA9-positive with a slight gradient in the previously shown patch prediction, it is crucial to highlight that the modeling framework’s predictions exhibit excellent alignment with patches devoid of CA9. This is evident in the results obtained for a patch from an Ovarian tumor tissue (0721) sample, illustrated in fig. S5. The scatter distribution of blood vessels in fig. S5(a) is well-captured by the framework, leading to a uniformly distributed oxygen pattern and corresponding absence of hypoxic regions, as depicted in Figures fig. S5(b) and fig. S5(d). Notably, this closely resembles the CA9 profile in fig. S5(c) with a very small validation error.

#### 3.4.2 Validation for whole sample

Detailed information at the patch level is valuable for examining microenvironments, but predicting hypoxia profiles for entire samples is equally crucial. Following the procedure outlined in section 2.12, we extend our predictions to whole samples, utilizing both IF and IHC scan types for testing. As illustrated in figs. 11(e) to 11(h) for an IF scan from a breast tumor tissue and fig. S7 for an IHC scan from a pancreas tumor, our predictions align well with the CA9 marker density. Similar to the observations at the patch level, our predictions cover CA9-positive cells and capture additional cells with a low hypoxia gradient.

It is essential to emphasize that identifying hypoxic cells is accurate; however, there are instances of false hypoxic regions being detected. This may be attributed to the coexistence of blood vessels and hypoxia or the non-availability of similar types of samples in the dataset.

## 4 Discussion

The well-established influence of hypoxia-driven alterations in the tumor microenvironment underscores the need for accurate assessments of oxygen and hypoxia distributions in effective cancer characterization and treatment design. We present a mechanistic framework applicable to various cancer types to predict hypoxic regions within the tumor microenvironment. This represents a crucial initial step towards a realistic model for hypoxia prediction and aiding in radiation dose estimation based on existing models [38, 48, 12, 49]. Understanding hypoxia is not only pivotal for radiation therapy efficacy but also essential for comprehending other hypoxia-induced phenomena, such as metabolic reprogramming, angiogenesis, tumor invasion and metastasis. In this study, we employ a simple mathematical approach capable of replicating biologically relevant hypotheses.

This data-driven yet mechanistic approach involves obtaining proxies for blood vessels, hypoxia, and cell nuclei (CD31, CA9, DAPI markers). We have formulated a mathematical model eq. (2.3) to derive oxygen distribution from blood vessel density and determine hypoxia profiles eq. (2.10). Optimization processes extract parameters by comparing model predictions with CA9 markers. The model is trained on tissue image patches, yielding parameter distributions. The formulated model eq. (2.3) is a linear PDE. We obtain similar results with models where oxygen production and uptake are limited, introducing nonlinearity (as mentioned in eq. (2.5), appendix B.1). However, due to the computational costs associated with solving the nonlinear PDE multiple times for optimization, and also to avoid overfitting and epistemic uncertainty resulting from extra parameters, we opt to stick with the linear model.

Assigning a class to each tissue patch proves beneficial to address the heterogeneity in blood vessels across different tissues and within the same tissue. We classify each patch based on its blood vessel architecture using the spatial entropy measure (refer to eq. (2.1)). This classification is integral to our prediction framework, where the first step involves identifying the patch class and obtaining the corresponding parameters. Additionally, a range of parameters is obtained for each classified patch, enabling the observation of different dynamics within the tissue.

Recognizing the limitations posed by the assumption of constant diffusion, production, and uptake of *O*_2_ molecules, we introduce a data-driven method to ascertain spatially varying coefficients. The method is based on the spatial entropy of patches from CD31 stained tissue images, summarizing the heterogeneity of blood vessels distributions, but the same approach can be adapted to other types of data such as hematoxylin and eosin staining of cell sections. Utilizing observed variations from trained and similar (classified) patches, we have employed a varying coefficient model (refer to eq. (2.12)) to quantify this heterogeneity. Through this process, we have deduced a reaction-diffusion PDE wherein the coefficients exhibit variability between the considered patches. This approach facilitates the use of tailored rates for different patch types, enabling predictions for the entire tissue sample using a simplified model.

Our modeling predictions perform well for both types of scans, IHC and IF, commonly used in practice for staining different markers. Specifically, IF is preferred due to computational efficiency, eliminating the need for image registration. Additionally, it utilizes the same tissue in experiments, simplifying both experimental and computational aspects as the cell structures remain consistent.

We have identified additional hypoxic regions beyond CA9 marker densities, possibly due to the 2D approximation of 3D vessels. To address this, we plan to enhance our model by incorporating the 3D geometry and architecture of blood vessels, moving beyond the 2D slices used in the current estimation. While 3D reconstruction, exemplified in the study [3], provides a more precise representation, its practicality in clinical settings is constrained by tissue availability. To overcome this limitation, we are developing a framework to quantify the approximation error when using 2D models, crucial for evaluating hypoxia profiles in scenarios where 3D reconstruction is limited.

While CA9 is often considered an intrinsic marker for hypoxia, studies such as [35, 19] for uterine cervix cancer and [24] for cervical carcinoma suggest a poor correlation between tumor oxygenation and CA9. This raises the need for exploring alternative markers or combinations of markers to improve the accuracy of hypoxia assessment. Additionally, the heterogeneity in oxygen consumption driven by cell metabolism is a crucial factor influencing hypoxia levels. Beyond the static assumptions and the 3D to 2D approximation, the dynamic state of cells may play a significant role in determining the hypoxia profile.

Predicting or estimating hypoxia without a time series dataset is valuable in various scenarios, especially given dynamic experiment challenges. Creating a profile based on the cell metabolic state and the functioning of blood vessels becomes essential, even if done less quantitatively, as attempted in our simulations for the diffusion coefficient, oxygen production, and uptake coefficients. Scores such as hypoxia scores (as seen in [51]) may need more information, as they often overlook spatial heterogeneity in both cells and hypoxia when applied to an entire image. Experimental investigations involving cell metabolic states and other markers to assess the state of blood vessels would be essential for validating the proposed profiling approach.

## 5 Conclusion

This study introduces an innovative data-driven yet mechanistic tool for characterizing tumor oxygenation, effectively capturing the intricate dynamics of the tumor microenvironment. The computational framework successfully reproduces these complex dynamics, as demonstrated in the context of three distinct tumor types: breast, ovarian, and pancreatic. The results not only facilitate localized predictions for small tissue regions but also offer a data-driven approach to extract spatially varying rates related to oxygen (such as diffusion coefficient, production, and uptake rate). This spatially resolved information is crucial for capturing and predicting spatial heterogeneity across larger tissue samples. Additionally, based on the derived oxygen-related rates, the study explores the potential to predict whether tumor vascularization is in a functioning or non-functioning state.

## Acknowledgment

We acknowledge financial support from Itmo Cancer on funds administered by INSERM (project MALMO) and a support by Mr Jean-Paul Baudecroux and The Big Brain Theory Program - Paris Brain Institute (ICM). We are grateful for the technical support of the Montpellier Rio Imaging (MRI) and animal (RAM) facilities. Histological analyses were performed thank to the “Réseau d’Histologie Expérimentale de Montpellier” (RHEM), a facility supported by the SIRIC Montpellier Cancer (Grant INCa_Inserm_DGOS_12553), the european regional development foundation and the Occitanie region (FEDER-FSE 2014-2020 Languedoc Roussillon), REACT-EU (Recovery Assistance for Cohesion and the Territories of Europe), IBiSA.

## A Supplementary Material

**Figure S1:**
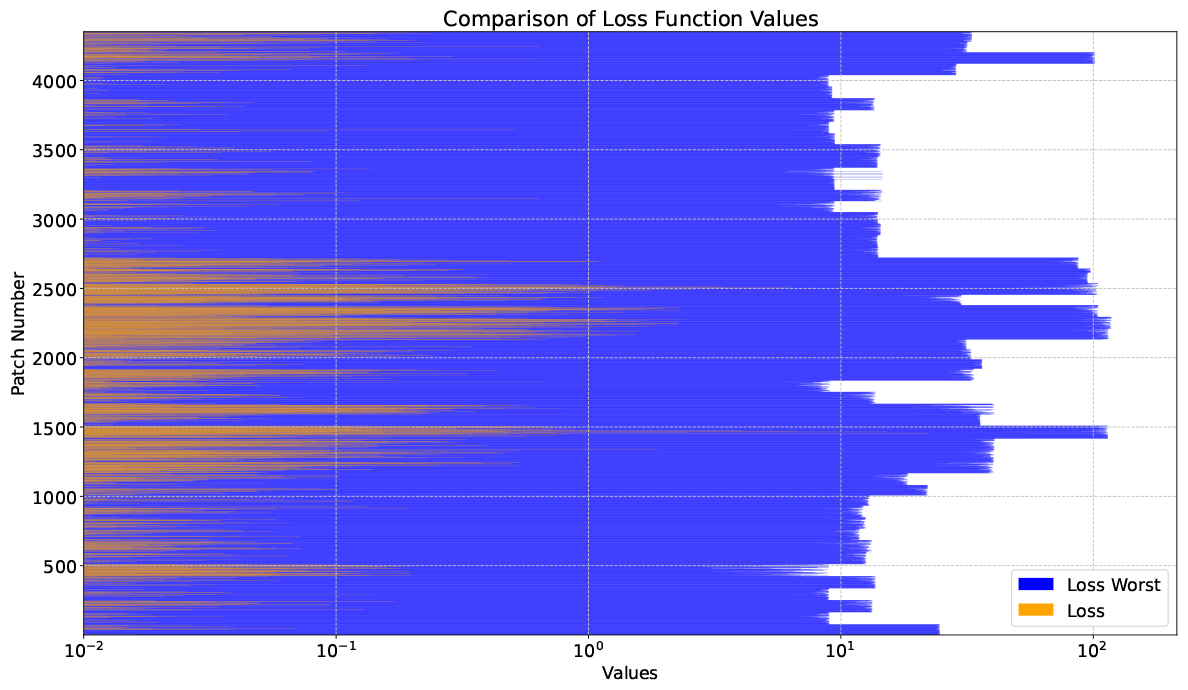
A comparison is made between the optimal loss function value from eq. (2.11) and its worst-case counterpart. The former is represented in orange, while the latter is depicted in blue. This comparison is conducted for each patch, observed along the y-axis, with corresponding values plotted on the x-axis. For each patch, the minimized values are significantly smaller than the worst values.

### A.1 IHC Results

**Figure S2:**
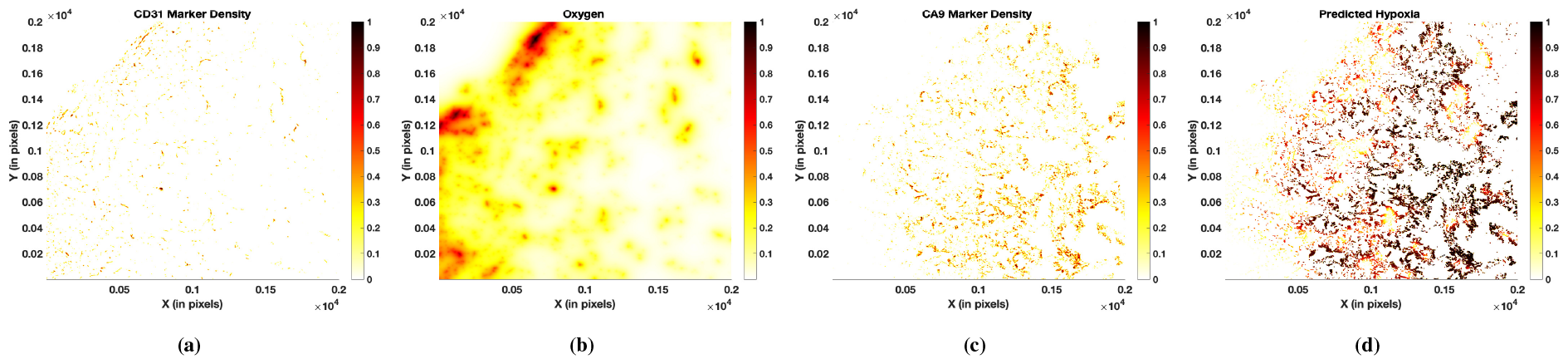
The data fitting also demonstrates effectiveness for the IHC scan. The presented plots are derived from a patch of pancreatic tumor tissue (1087) image and an IHC scan. Plots (a) and (c) showcase the densities of blood vessels and CA9, obtained through the procedure outlined in section 2.4. Meanwhile, plots (b) and (d) illustrates the oxygen and hypoxia profile for the optimized set of parameters estimated through the optimization process detailed in section 2.10. In this case, the loss function value = 0.04, while the worst case value = 113.95.

#### A.1.1 Patchwise

The patchwise validation results for ovarian (0721) and pancreatic (1087) tumors, obtained from IHC type staining, are shown in Figures S5 and S6.

#### A.1.2 Whole sample

The whole sample validation for a pancreatic tumor (1905), obtained from an IHC-type scan, is depicted in Figure S7.

**Figure S3:**
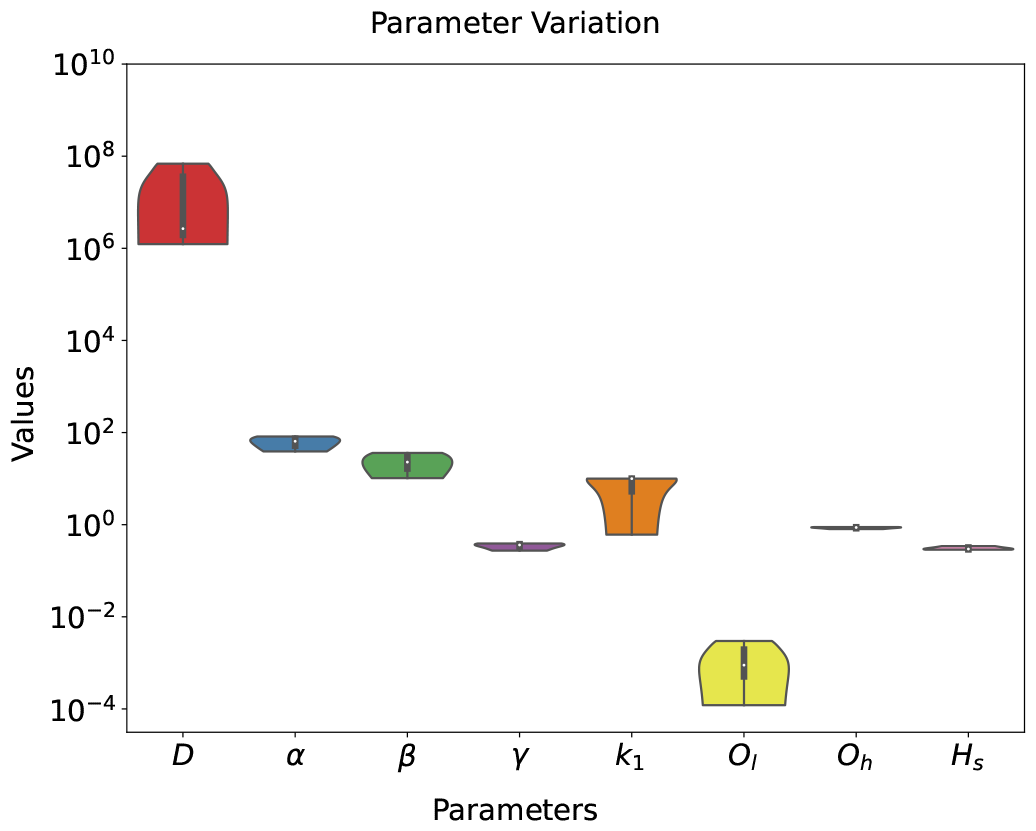
Similar reduced variability of the involved coefficients is observed for another cancer type. This figure illustrates the obtained interval of parameters based on blood vessel architecture for a pancreatic tumor scan of IHC type from left to right: diffusion coefficient *D, O*_2_ uptake and production rates (*α, β*), *O*_*l*_, *O*_*h*_, and heterogeneity score for the trained samples. The reduced variability across different coefficients suggests consistent trends in tumor oxygenation possibilities.

**Figure S4:**
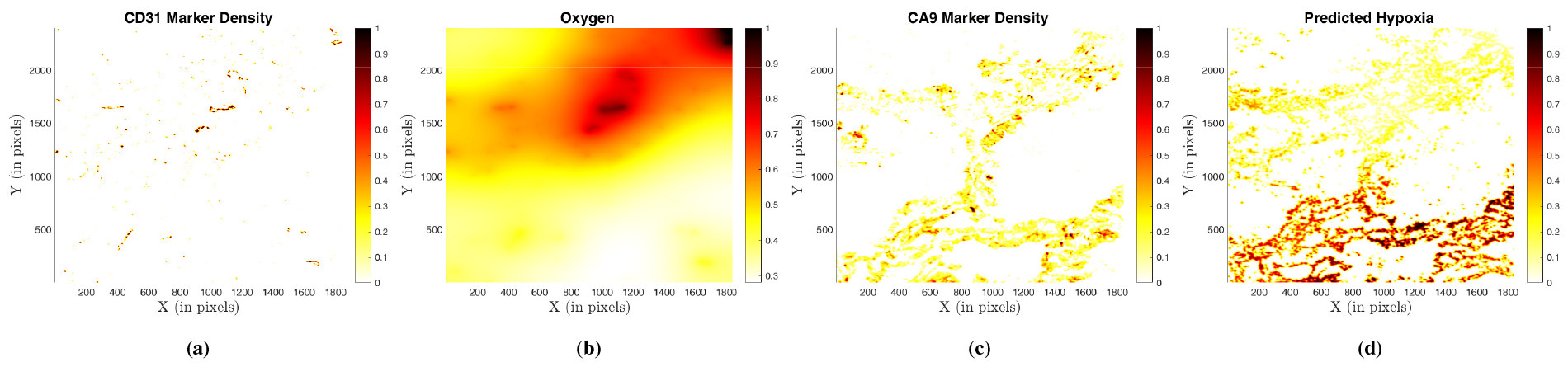
The predictions align well for a pancreatic tumor sample (1087) of IFC type scan, as observed from the visual representation of the predicted hypoxia (*h*) using eq. (2.3) and (2.10). Quantitatively, the validation error is 0.5163.

**Figure S5:**
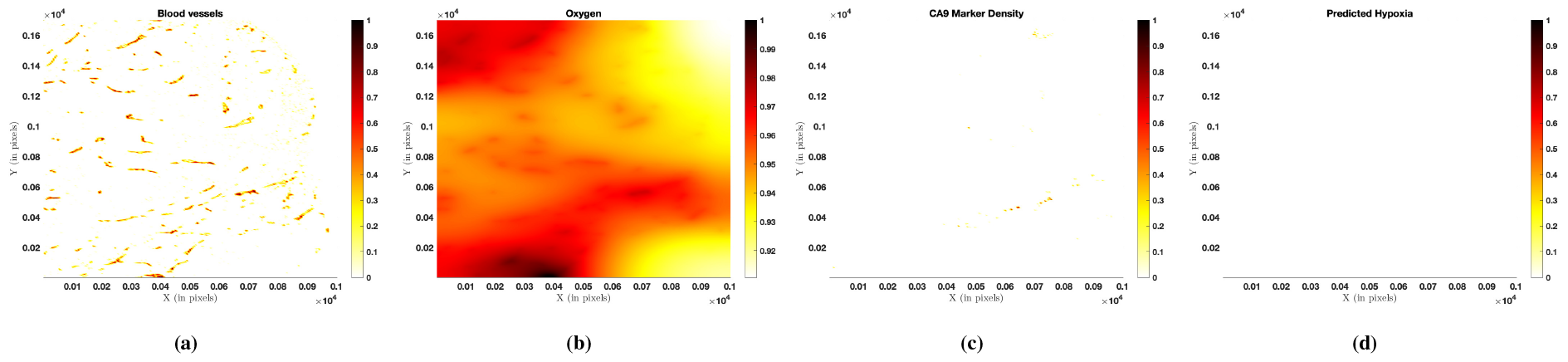
Similar to the results showcased for IF scans in the main text, the model simulations exhibit effectiveness for IHC scans. The predictions for a patch of ovarian tumor sample (0721) from an IHC-type scan show good agreement. This is evident in the visual representation of the predicted hypoxia (*h*) using eq. (2.3) and (2.10). The quantitative assessment further supports the alignment, with a validation error 0.0036.

## B Other forms of the model

### B.1 Model II: Limited uptake model

To observe the limitation in the uptake term, we propose a logistic type uptake as

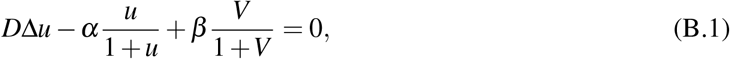

**Figure S6:**
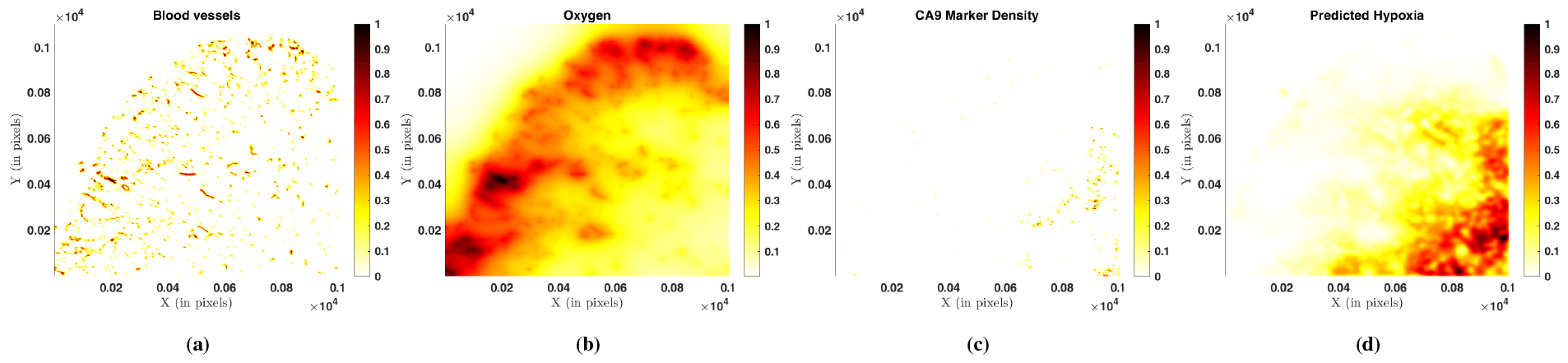
For a patch of the pancreatic tumor sample (1087) from an IHC-type scan, the predicted hypoxia closely resembles the CA9 density, with a validation error of 1.3893. However, additional hypoxic regions are observed beyond the CA9 marker densities in this case.

**Figure S7:**
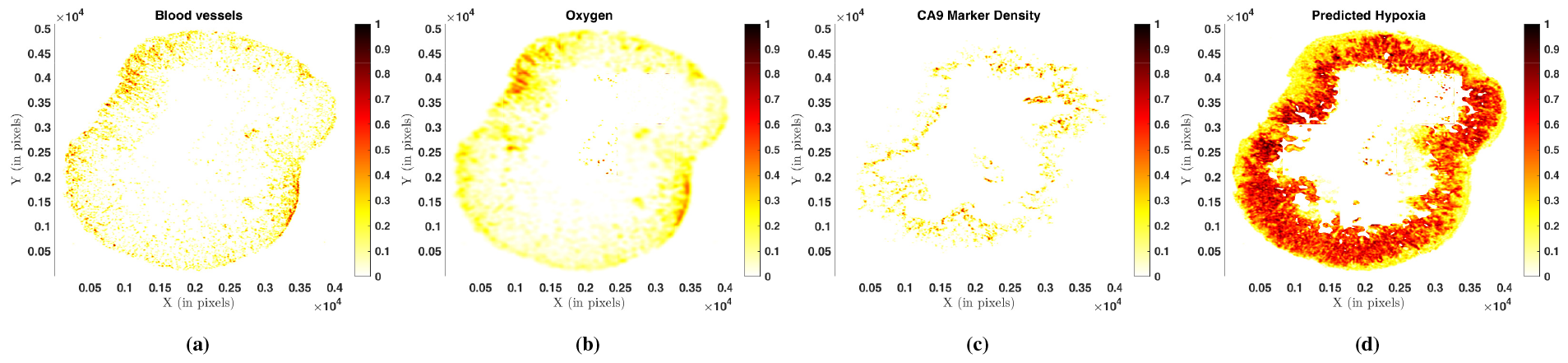
The prediction and validation for the entire tissue sample also show successful results for IHC-type staining. This is evident in the case of a pancreatic tumor (1905), where the model predictions, shown in plots (b) and (d) for *O*_2_ and hypoxia profiles, respectively, align well with the CA9 data presented in (c). The model captures additional regions of hypoxia, resulting in a validation error of 6.6968

again with the boundary conditions: ∇*u n* = 0. This formulation yields a nonlinear model. However, similar results have been observed, as shown in earlier sections.

### B.2 Data

The experimentally generated images will be shared.

### Codes

The codes will be available publicly upon publication at: https://github.com/its-Pa1/2D_Hypoxia_Modeling

## Footnotes

1 The abbreviation “CD31” refers to “Cluster of Differentiation 31”, a biomarker commonly utilized for identifying blood vessels, particularly in tumor tissues, as detailed in the next paragraph.

2 In this study, we sometimes use the terms hypoxia and degree/measure of hypoxia interchangeably. Hypoxia is a state/condition of tissue environment represented by CA9 marker density in this work.

3 MATLAB Version: 9.13.0.2126072 (R2022b), The MathWorks Inc., Natick, Massachusetts, United States

4 ZEISS ZEN (blue edition), Version 3.6.095.01000, Carl Zeiss Microscopy GmbH, Germany

5 The Gaussian filter *G* for image *I* in 2D is defined as *I ∗ G*, where 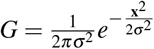, with *σ* being the standard deviation with zero mean applied to each point **x** = (*x, y*) on a 2D grid.

6 The density 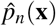 is estimated 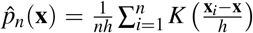 with *K*(**x**) being a Gaussian kernel function given by 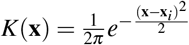,*h* being the smoothing bandwidth, **x**_*i*_ be the observed data point, **x** is the value where kernel function is computed and *n* being the number of data points involved.

7 In both methods, a point (in Euclidean space) is represented by the pixel coordinates of an image.

8 Throughout this classification, the term “area” is used to denote the number of pixels in a connected component, which remains independent of the actual pixel dimensions in each image. When applying eq. (2.1), we require the proportion of such connected components, making it independent of the actual pixel length and breadth. Similarly, distances are calculated based on pixel coordinates, so the ratio of distances involved in eq. (2.1) is also independent of actual pixel dimensions. This approach ensures that the comparison of *H*_*s*_ between different image patches is meaningful.

9 A sharp transition between states, in this case, from normoxic to (severe) hypoxic state or vice-versa

## Notes

### Competing Interest Statement

The authors have declared no competing interest.

## References

[1] W. Al Tameemi, T. P. Dale, R. M. K. Al-Jumaily, and N. R. Forsyth. “Hypoxia-Modified Cancer Cell Metabolism”. In: Frontiers in Cell and Developmental Biology 7 (2019). DOI: 10.3389/fcell.2019.00004.

[2] P. M. Altrock, L. L. Liu, and F. Michor. “The mathematics of cancer: Integrating quantitative models”. In: Nature Reviews Cancer 15.12 (2015), pp. 730–745. DOI: 10.1038/nrc4029.

[3] J. Arslan, M. Ounissi, H. Luo, M. Lacroix, P. Dupré, P. Kumar, A. Hodgkinson, S. Dandou, R. Larive, C. Pignodel, L. L. Cam, O. Radulescu, and D. Racoceanu. “Efficient 3D reconstruction of whole slide images in melanoma”. In: Medical Imaging 2023: Digital and Computational Pathology. Vol. 12471. SPIE, 2023, 124711S. DOI: 10.1117/12.2657473.

[4] B. Bedogni and M. B. Powell. “Hypoxia, melanocytes and melanoma - survival and tumor development in the permissive microenvironment of the skin”. In: Pigment Cell & Melanoma Research 22.2 (2009), pp. 166–174. DOI: 10.1111/j.1755-148X.2009.00553.x.

[5] T. van den Beucken, M. Koritzinsky, H. Niessen, L. Dubois, K. Savelkouls, H. Mujcic, B. Jutten, J. Kopacek, S. Pastorekova, A. J. van der Kogel, et al. “Hypoxia-induced expression of carbonic anhydrase 9 is dependent on the unfolded protein response”. In: Journal of Biological Chemistry 284.36 (2009), pp. 24204–24212. DOI: 10.1074/jbc.M109.006510.

[6] D. J. Carlson, P. J. Keall, B. W. Loo, Z. J. Chen, and J. M. Brown. “Hypofractionation Results in Reduced Tumor Cell Kill Compared to Conventional Fractionation for Tumors With Regions of Hypoxia”. In: International Journal of Radiation Oncology, Biology, Physics 79.4 (2011), pp. 1188–1195. DOI: 10.1016/j.ijrobp.2010.10.007.

[7] M. A. Cavadas, L. K. Nguyen, and A. Cheong. “Hypoxia-inducible factor (HIF) network: Insights from mathematical models”. In: Cell Communication and Signaling 11.1 (2013). DOI: 10.1186/1478-811X-11-42.

[8] C. Claramunt. “A spatial form of diversity”. In: Spatial Information Theory: International Conference, COSIT 2005, Ellicottville, NY, USA, September 14-18, 2005. Proceedings 7. Springer. 2005, pp. 218–231. DOI: 10.1007/11556114_14.

[9] P. E. Colombo, S. du Manoir, B. Orsetti, R. Bras-Gonçalves, M. B. Lambros, A. MacKay, T. T. Nguyen, F. Boissière, D. Pourquier, F. Bibeau, J. S. Reis-Filho, and C. Theillet. “Ovarian carcinoma patient derived xenografts reproduce their tumor of origin and preserve an oligoclonal structure”. In: Oncotarget 6.29 (2015), pp. 28327–28340. DOI: 10.18632/oncotarget.5069.

[10] S. D’Aguanno, F. Mallone, M. Marenco, D. Del Bufalo, and A. Moramarco. “Hypoxia-dependent drivers of melanoma progression”. In: Journal of Experimental and Clinical Cancer Research 40.1 (2021), pp. 1–32. DOI: 10.1186/s13046-021-01926-6.

[11] A. Daşu, I. Toma-Daşu, and M. Karlsson. “Theoretical simulation of tumour oxygenation and results from acute and chronic hypoxia”. In: Physics in Medicine and Biology 48.17 (2003), pp. 2829–2842. DOI: 10.1088/0031-9155/48/17/307.

[12] A. Daşu, I. Toma-Daşu, and M. Karlsson. “The effects of hypoxia on the theoretical modelling of tumour control probability”. In: Acta Oncologica 44.6 (2005), pp. 563–571. DOI: 10.1080/02841860500244435.

[13] B. Gallez, M. A. Neveu, P. Danhier, and B. F. Jordan. “Manipulation of tumor oxygenation and radiosensitivity through modification of cell respiration. A critical review of approaches and imaging biomarkers for therapeutic guidance”. In: Biochimica et Biophysica Acta - Bioenergetics 1858.8 (2017), pp. 700–711. DOI: 10.1016/j.bbabio.2017.01.002.

[14] A. Ginouvès, K. Ilc, N. Macías, J. Pouysségur, and E. Berra. “PHDs overactivation during chronic hypoxia “desensitizes” HIFα and protects cells from necrosis”. In: Proceedings of the National Academy of Sciences of the United States of America 105.12 (2008), pp. 4745–4750. DOI: 10.1073/pnas.0705680105.

[15] D. Goldman. “Theoretical Models of Microvascular Oxygen Transport to Tissue”. In: Microcirculation 15.8 (2008), pp. 795–811. DOI: 10.1080/10739680801938289. arXiv: NIHMS150003.

[16] D. R. Grimes, P. Kannan, D. R. Warren, B. Markelc, R. Bates, R. Muschel, and M. Partridge. “Estimating oxygen distribution from vasculature in three-dimensional tumour tissue”. In: Journal of The Royal Society Interface 13.119 (2016), p. 20160362. DOI: 10.1098/rsif.2016.0362.

[17] T. M. Grzywa, W. Paskal, and P. K. Włodarski. “Intratumor and Intertumor Heterogeneity in Melanoma”. In: Translational Oncology 10.6 (2017), pp. 956–975. DOI: 10.1016/j.tranon.2017.09.007.

[18] A.L. Harris. “Hypoxia - A key regulatory factor in tumour growth”. In: Nature Reviews Cancer 2.1 (2002), pp. 38–47. DOI: 10.1038/nrc704.

[19] D. Hedley, M. Pintilie, J. Woo, A. Morrison, D. Birle, A. Fyles, M. Milosevic, and R. Hill. “Carbonic Anhydrase IX Expression, Hypoxia, and Prognosis in Patients with Uterine Cervical Carcinomas”. In: Clinical Cancer Research 9.15 (2003), pp. 5666–5674. eprint: https://aacrjournals.org/clincancerres/article-pdf/9/15/5666/2085276/zdf01503005666.pdf.

[20] K. Hirota and G. L. Semenza. “Regulation of angiogenesis by hypoxia-inducible factor 1”. In: Critical Reviews in Oncology/Hematology 59.1 (2006), pp. 15–26. DOI: 10.1016/j.critrevonc.2005.12.003.

[21] A. Hodgkinson, D. Trucu, M. Lacroix, L. Le Cam, and O. Radulescu. “Computational Model of Heterogeneity in Melanoma: Designing Therapies and Predicting Outcomes”. In: Frontiers in oncology (2022), p. 1245. DOI: 10.3389/fonc.2022.857572.

[22] M. R. Horsman and A. J. van der Kogel. “Therapeutic approaches to tumour hypoxia”. In: Basic clinical radiobiology 20.40 (2009), p. 233. DOI: 10.1201/b15450-20.

[23] R. Hsu and T. W. Secomb. “A Green’s function method for analysis of oxygen delivery to tissue by microvascular networks”. In: Mathematical Biosciences 96.1 (1989), pp. 61–78. DOI: 10.1016/0025-5564(89)90083-7.

[24] V. V. Iakovlev, M. Pintilie, A. Morrison, A. W. Fyles, R. P. Hill, and D. W. Hedley. “Effect of distributional heterogeneity on the analysis of tumor hypoxia based on carbonic anhydrase IX”. In: Laboratory Investigation 87.12 (2007), pp. 1206–1217. DOI: 10.1038/labinvest.3700680.

[25] P. Jorjani and S. Ozturk. “Effects of Cell Density and Temperature Different Mammalian Cell Lines”. In: Biotechnology and bioengineering 64.3 (1999), pp. 349–356. DOI: 10.1002/(SICI)1097-0290(19990805)64:3<349::AID-BIT11>3.0.CO; 2-V.

[26] J. Kapuscinski. “DAPI: a DNA-specific fluorescent probe”. In: Biotechnic & histochemistry 70.5 (1995), pp. 220–233. DOI: 10.3109/10520299509108199.

[27] C. J. Kelly and M. Brady. “A model to simulate tumour oxygenation and dynamic [18F]-Fmiso PET data”. In: Physics in Medicine and Biology 51.22 (2006), pp. 5859–5873. DOI: 10.1088/0031-9155/51/22/009.

[28] H. Kempf, M. Bleicher, and M. Meyer-Hermann. “Spatio-temporal dynamics of hypoxia during radiotherapy”. In: PLoS One 10.8 (2015), e0133357. DOI: 10.1371/journal.pone.0133357.

[29] J. Korbecki, D. Simińska, M. Gąssowska-Dobrowolska, J. Listos, I. Gutowska, D. Chlubek, and I. Baranowska-Bosiacka. “Chronic and cycling hypoxia: drivers of cancer chronic inflammation through HIF-1 and NF-κB activation: a review of the molecular mechanisms”. In: International Journal of Molecular Sciences 22.19 (2021), p. 10701. DOI: 10.3390/ijms221910701.

[30] A. Krogh. “The number and distribution of capillaries in muscles with calculations of the oxygen pressure head necessary for supplying the tissue”. In: The Journal of Physiology 52.6 (1919), pp. 409–415. DOI: 10.1113/jphysiol.1919.sp001839.

[31] P. Kumar and C. Surulescu. “A Flux-Limited Model for Glioma Patterning with Hypoxia-Induced Angiogenesis”. In: Symmetry 12.11 (2020). DOI: 10.3390/sym12111870.

[32] A. Lin and A. Maity. “Molecular Pathways: A Novel Approach to Targeting Hypoxia and Improving Radiotherapy Efficacy via Reduction in Oxygen Demand”. In: Clinical Cancer Research 21.9 (2015), pp. 1995–2000. DOI: 10.1158/1078-0432.CCR-14-0858.

[33] N. J. Mabjeesh and S. Amir. “Hypoxia-inducible factor (HIF) in human tumorigenesis”. In: Histology and Histopathology 22.4-6 (2007), pp. 559–572. DOI: 10.14670/HH-22.559.

[34] S. du Manoir, B. Orsetti, R. Bras-Gonçalves, T. T. Nguyen, L. Lasorsa, F. Boissière, B. Massemin, P. E. Colombo, F. Bibeau, W. Jacot, and C. Theillet. “Breast tumor PDXs are genetically plastic and correspond to a subset of aggressive cancers prone to relapse”. In: Molecular Oncology 8.2 (2014), pp. 431–443. DOI: 10.1016/j.molonc.2013.11.010.

[35] A. Mayer, M. Höckel, and P. Vaupel. “Carbonic anhydrase IX expression end tumor oxygenation status do not correlate at the microregional level in locally advanced cancers of the uterine cervix”. In: Clinical Cancer Research 11.20 (2005), pp. 7220–7225. DOI: 10.1158/1078-0432.CCR-05-0869.

[36] S. R. McKeown. “Defining normoxia, physoxia and hypoxia in tumours - Implications for treatment response”. In: British Journal of Radiology 87.1035 (2014), pp. 1–12. DOI: 10.1259/bjr.20130676.

[37] L. K. Nguyen, M. A. S. Cavadas, C. C. Scholz, S. F. Fitzpatrick, U. Bruning, E. P. Cummins, M. M. Tambuwala, M. C. Manresa, B. N. Kholodenko, C. T. Taylor, and A. Cheong. “A dynamic model of the hypoxia-inducible factor 1a (HIF-1a) network”. In: Journal of Cell Science 128.2 (2015), p. 422. DOI: 10.1242/jcs.167304.

[38] G. Powathil, M. Kohandel, M. Milosevic, and S. Sivaloganathan. “Modeling the spatial distribution of chronic tumor hypoxia: Implications for experimental and clinical studies”. In: Computational and Mathematical Methods in Medicine 2012 (2012). DOI: 10.1155/2012/410602.

[39] M. P. Pusztaszeri, W. Seelentag, and F. T. Bosman. “Immunohistochemical Expression of Endothelial Markers CD31, CD34, von Willebrand Factor, and Fli-1 in Normal Human Tissues”. In: Journal of Histochemistry & Cytochemistry 54.4 (2006). PMID: 16234507, pp. 385–395. DOI: 10.1369/jhc.4A6514.2005.

[40] A. A. Qutub and A. S. Popel. “A computational model of intracellular oxygen sensing by hypoxia-inducible factor HIF1α”. In: Journal of Cell Science 119.16 (2006), pp. 3467–3480. DOI: 10.1242/jcs.03087.

[41] K. Saxena and M. K. Jolly. “Acute vs. Chronic vs. cyclic hypoxia: Their differential dynamics, molecular mechanisms, and effects on tumor progression”. In: Biomolecules 9.8 (2019), pp. 1–27. DOI: 10.3390/biom9080339.

[42] T. W. Secomb, R. Hsu, M. W. Dewhirst, B. Klitzman, and J. F. Gross. “Analysis of oxygen transport to tumor tissue by microvascular networks”. In: International Journal of Radiation Oncology, Biology, Physics 25.3 (1993), pp. 481–489. DOI: 10.1016/0360-3016(93)90070-C.

[43] T. W. Secomb, R. Hsu, E. T. Ong, J. F. Gross, and M. W. Dewhirst. “Analysis of the effects of oxygen supply and demand on hypoxic fraction in tumors”. In: Acta Oncologica 34.3 (1995), pp. 313–316. DOI: 10.3109/02841869509093981.

[44] T. W. Secomb, R. Hsu, E. Y. Park, and M. W. Dewhirst. “Green’s function methods for analysis of oxygen delivery to tissue by microvascular networks”. In: Annals of Biomedical Engineering 32.11 (2004), pp. 1519–1529. DOI: 10.1114/B:ABME.0000049036.08817.44.

[45] A. C. Skeldon, G. Chaffey, D. J. Lloyd, V. Mohan, D. A. Bradley, and A. Nisbet. “Modelling and detecting tumour oxygenation levels”. In: PLoS ONE 7.6 (2012), pp. 1–17. DOI: 10.1371/journal.pone.0038597.

[46] J. H. Suh, B. Stea, A. Nabid, J. J. Kresl, A. Fortin, J. P. Mercier, N. Senzer, E. L. Chang, A. P. Boyd, P. J. Cagnoni, and E. Shaw. “Phase III study of efaproxiral as an adjunct to whole-brain radiation therapy for brain metastases”. In: Journal of Clinical Oncology 24.1 (2006), pp. 106–114. DOI: 10.1200/JCO.2004.00.1768.

[47] B. I. Tarnowski, F. G. Spinale, and J. H. Nicholson. “DAPI as a useful stain for nuclear quantitation”. In: Biotechnic & histochemistry 66.6 (1991), pp. 296–302. DOI: 10.3109/10520299109109990.

[48] B. Titz and R. Jeraj. “An imaging-based tumour growth and treatment response model: Investigating the effect of tumour oxygenation on radiation therapy response”. In: Physics in Medicine and Biology 53.17 (2008), pp. 4471–4488. DOI: 10.1088/0031-9155/53/17/001.

[49] I. Toma-Dasu and A. Dasu. “Modelling tumour oxygenation, reoxygenation and implications on treatment outcome”. In: Computational and Mathematical Methods in Medicine 2013 (2013). DOI: 10.1155/2013/141087.

[50] K. J. Turner, J. P. Crew, C. C. Wykoff, P. H. Watson, R. Poulsom, J. Pastorek, P. J. Ratcliffe, D. Cranston, and A. L. Harris. “The hypoxia-inducible genes VEGF and CA9 are differentially regulated in superficial vs invasive bladder cancer”. In: British Journal of Cancer 86.8 (2002), pp. 1276–1282. DOI: 10.1038/sj.bjc.6600215.

[51] H. W. Van Laarhoven, J. H. Kaanders, J. Lok, W. J. Peeters, P. F. Rijken, B. Wiering, T. J. Ruers, C. J. Punt, A. Heerschap, and A. J. Van Der Kogel. “Hypoxia in relation to vasculature and proliferation in liver metastases in patients with colorectal cancer”. In: International Journal of Radiation Oncology Biology Physics 64.2 (2006), pp. 473–482. DOI: 10.1016/j.ijrobp.2005.07.982.

[52] P. Vaupel, A. B. Flood, and H. M. Swartz. “Oxygenation Status of Malignant Tumors vs. Normal Tissues: Critical Evaluation and Updated Data Source Based on Direct Measurements with pO2 Microsensors”. In: Applied Magnetic Resonance 52.10 (2021), pp. 1451–1479. DOI: 10.1007/s00723-021-01383-6.

[53] P. Vaupel and A. Mayer. “Hypoxia in cancer: Significance and impact on clinical outcome”. In: Cancer and Metastasis Reviews 26.2 (2007), pp. 225–239. DOI: 10.1007/s10555-007-9055-1.

[54] M. Welter, T. Fredrich, H. Rinneberg, and H. Rieger. “Computational model for tumor oxygenation applied to clinical data on breast tumor hemoglobin concentrations suggests vascular dilatation and compression”. In: PLoS ONE 11.8 (2016), pp. 1–42. DOI: 10.1371/journal.pone.0161267.

[55] D. S. Widmer, K. S. Hoek, P. F. Cheng, O. M. Eichhoff, T. Biedermann, M. I. Raaijmakers, S. Hemmi, R. Dummer, and M. P. Levesque. “Hypoxia contributes to melanoma heterogeneity by triggering HIF1α-dependent phenotype switching”. In: Journal of Investigative Dermatology 133.10 (2013), pp. 2436–2443. DOI: 10.1038/jid.2013.115.

